# AAclust: *k*-optimized clustering for selecting redundancy-reduced sets of amino acid scales

**DOI:** 10.1101/2024.02.04.578800

**Authors:** Stephan Breimann, Dmitrij Frishman

## Abstract

**Summary:** Amino acid scales are crucial for sequence-based protein prediction tasks, yet no gold standard scale set or simple scale selection methods exist. We developed AAclust, a wrapper for clustering models that require a pre-defined number of clusters *k*, such as *k-*means. AAclust obtains redundancy-reduced scale sets by clustering and selecting one representative scale per cluster, where *k* can either be optimized by AAclust or defined by the user. The utility of AAclust scale selections was assessed by applying machine learning models to 24 protein benchmark datasets. We found that top-performing scale sets were different for each benchmark dataset and significantly outperformed scale sets used in previous studies. Notably, model performance showed a strong positive correlation with the scale set size. AAclust enables a systematic optimization of scale-based feature engineering in machine learning applications.

**Availability and implementation:** The AAclust algorithm is part of AAanalysis, a Python-based framework for interpretable sequence-based protein prediction, which will be made freely accessible in a forthcoming publication.

**Contact:** Stephan Breimann and Dmitrij Frishman

**Supplementary information:** Further details on methods and results are provided in Supplementary Material.

## 1 Introduction

Starting with the influential works of Sneath, Bigelow, and Zimmerman [1–3] in the 1960s, amino acids have been described by numerical indices or scales reflecting their physicochemical properties, such as volume, polarity, or charge. The AAindex database [4] currently contains 566 experimentally measured or computationally derived indices published in 149 studies over 6 decades. However, this dataset is highly redundant—for example, over 120 scales are dedicated to polarity and α-helix propensity. While subsets of AAindex [5–8] are commonly used in sequence-based machine learning applications, typically selected based on heuristic criteria, a universally accepted ‘gold-standard’ scale set has so far been lacking.

Redundancy increases the data dimensionality, leading potentially to a bias towards repetitive information and overfitting in machine learning applications [9]. Reducing such redundancies can improve the efficiency and performance of algorithms, while also enhancing their general interpretability [10]. Redundancy reduction is a common step in a variety of bioinformatics applications, such as summarizing gene ontology [11] term lists (e.g., via REVIGO [12]), or creating redundancy-reduced protein sequence sets (e.g., via CD-HIT [13]). These methods are designed to cluster data based on similarity measures, such as semantic or sequence similarity, and then select a single representative per cluster. In this vein, we introduce AAclust, a clustering framework leveraging Pearson correlation as a similarity measure to select redundancy-reduced amino acid scale sets. Using machine learning models, we assessed the performance of AAclust scale selections against ‘gold standard’, randomly selected, and principal component-based scales.

## 2 Materials and methods

### 2.1 Dataset collation

#### Amino acid scales

We assembled a set of 586 amino acid scales (SCALES, **Supplementary Table 1**), by first obtaining 553 scales from AAindex that do not contain missing values. We included 21 further scales regarding accessible surface area (ASA) from Lins et al. [14] and 12 hydrophobicity scales from Koehler et al. [15] because of their relevance for protein folding [16] and backbone dynamics [17]. Each scale was min-max normalized to the range of [0, 1].

### Scale sets for benchmarking

Scale sets selected by AAclust were compared against three groups of baseline scale sets:

- **standard**: Three ‘gold standard’ sets, two from previous studies comprising 7 [18,19] and 12 [20] scales (**Supplementary Table 1**), and all 586 scales from SCALES.
- **pc-based**: All scales from SCALES transformed using principal component (PC) analysis into 20 principal components, each serving as a single scale, named P1 to P20.
- **random:** Subsets of varying size assembled by randomly sampling scales from SCALES.

### Datasets of protein sequences

We collated 12 protein sequence datasets (**Table S1, Supplementary Table 2**) from previous studies targeting distinct binary classification tasks: 6 datasets were used to predict entire protein sequence properties, while the other 6 served to predict residue properties in specific sequence positions. These groups are referred to as ‘sequence prediction’ and ‘residue prediction’ dataset, respectively, with individual dataset names based on the prediction task (‘SEQ_*dataset_name*’ or ‘AA_*dataset_name*’). For residue predictions, we used three amino acid window sizes (*n*=5, 9, 13), resulting in a total of 24 benchmark datasets. Each benchmark dataset was balanced by random sampling 400 data points (proteins or residues) per class.

### 2.2 AAclust: *k*-optimized clustering

AAclust is a clustering wrapper [21,22] framework extending clustering models that require a pre-defined number of clusters *k*, such as *k-*means [23], thereby eliminating the need to specify *k* in advance. It automatically partitions scale sets into *k* clusters by maximizing the within-cluster Pearson correlation to surpass a user-defined minimum threshold *min_th*. Two alternative quality measures are employed: the minimum pairwise Pearson correlation among all cluster members (‘*min_cor*_*all*_’) or the minimum Pearson correlation between the cluster center and all cluster members (‘*min_cor*_*center*_’). The minimum correlation across all clusters can be maximized using either *min_cor*_*all*_ or *min_cor*_*center*_.

Optimizing *k* in a three-step procedure (**Fig. 1a**), AAclust first estimates the lower bound of *k*, then refines it through recursive clustering (using *min_cor*_*all*_ or *min_cor*_*center*_), and optionally merges smaller clusters into larger ones based on Pearson correlation or Euclidean distance. AAclust is controlled by three parameters:

**Fig. 1.**
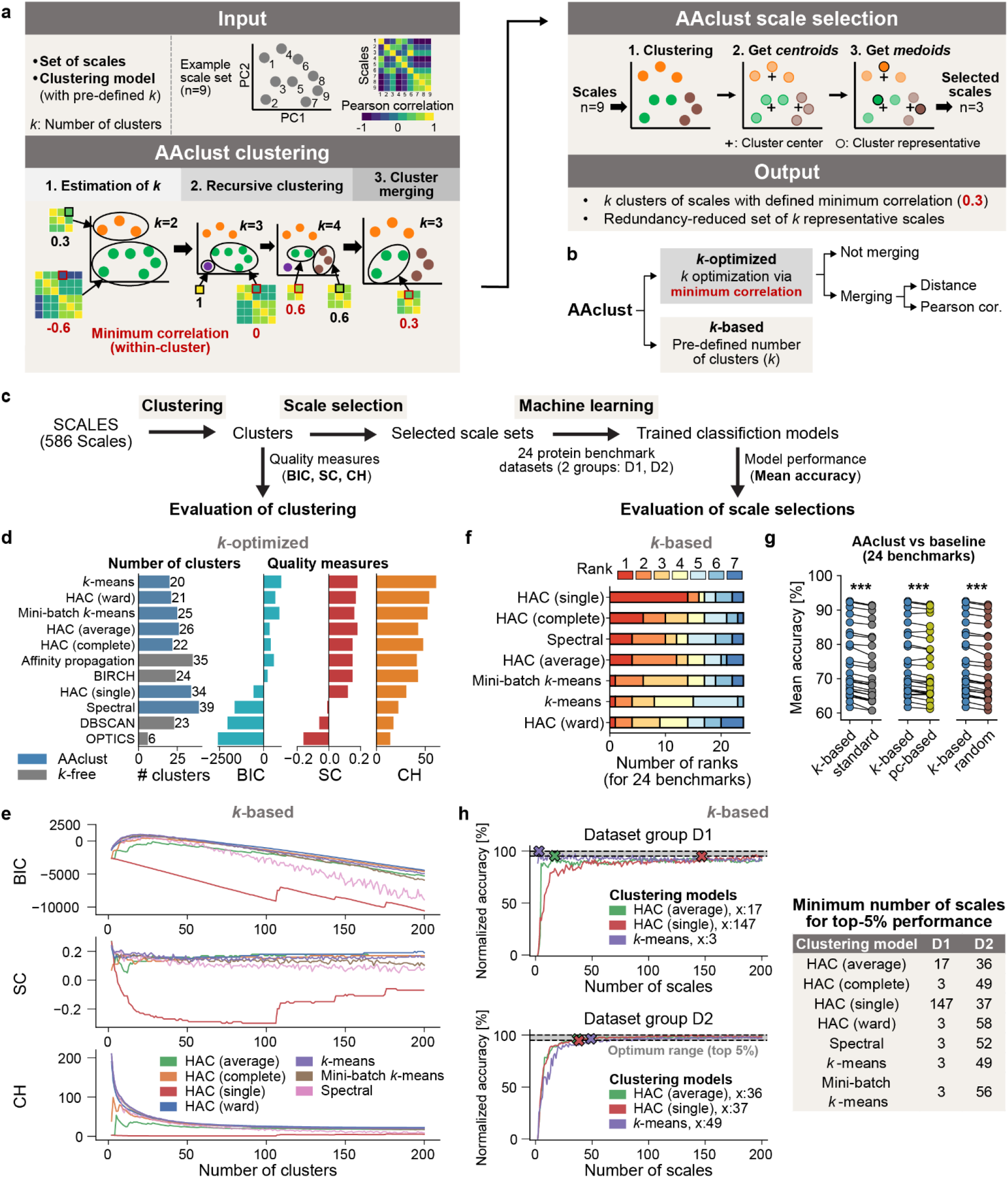
AAclust algorithm, workflow, and evaluation. **a-c**, Overview of the AAclust algorithm, settings, and evaluation workflow. **d-e**, Evaluation of AAclust clustering using Bayesian information criterion (BIC), silhouette coefficient (SC), the Calinski Harabasz score (CH). **d**, Comparison of the best *k*-optimized AAclust approaches against *k-*free clustering models. **e**, Relation between the number of clusters and the quality measures for *k*-based AAclust approaches. **f-g**, Evaluation of AAclust scale selection. **f**, The number of ranks of best-performing clustering models for *k*-based AAclust approaches. **g**, Comparison of best-performing (for each benchmark dataset) *k*-based AAclust approaches against baseline scale sets: ‘standard’ (gray), ‘pc-based’ (yellow), and ‘random’ (brown). Differences were tested by paired Wilcoxon signed-rank test, Benjamin Hochberg correction (*P<0.05,**P<0.01, ***P<0.001). **h**, Relation between the number of scales and the aggregated prediction performance for *k*-based AAclust approaches. ‘x: *n*’ indicates the minimum number of scales *n* for a top-5% performance.

- ***min_th***: Sets the Pearson correlation threshold (between 0 and 1, default *min_th*=0.3) to define the minimum correlation for all clusters.
- **Center**: Determines whether *min_th* applies to the cluster center (true) or all cluster members (false), using either ‘*min_cor*_*center*_’ or ‘*min_cor*_*all*_’, respectively.
- **Merge**: Enables (true) or disables (false) the optional merging step.To obtain redundancy-reduced scale sets, AAclust selects one representative scale per cluster, closest to its center. Alternatively, users can set *k* to a pre-defined number of scales. Both methods are referred to as ‘*k*-optimized’ and ‘*k*-based’ approaches (**Fig. 1b**), respectively.

### 2.3 Quality measures of clustering

To evaluate cluster quality (**Fig. 1c**), we clustered all scales from SCALES by seven clustering models used with the *k*-optimized AAclust approach and three clustering models that do not require a pre-specified *k*, referred to as ‘*k-*free’ clustering models (**Table S2**). Three commonly used clustering quality measures [24–26] were employed: silhouette coefficient (SC) [27,28], Calinski Harabasz score (CH) [29], and Bayesian information criterion (BIC) [30] (**Table S3**).

### 2.4 Evaluation procedure for scale selections

To assess AAclust scale selections (**Fig. 1c**), we compared them to ‘standard’, ‘pc-based’, and ‘random’ scale sets. AAclust was employed with seven clustering models, testing different *k*-optimized AAclust settings and assessing various scale set ranges for *k*-based approaches. Each scale set served as a feature set for 24 benchmark datasets using three machine learning models with default settings: random forest, support vector machine, and logistic regression. Model performance was measured by accuracy (ACC) using five-fold cross validation. To minimize model-dependent bias, we averaged accuracy across all folds and models (‘mean accuracy’), using it as the quality measure for each scale set.

## 3 Results

### 3.1 Evaluation of clustering

We comprehensively evaluated the quality of clusters derived from the entire SCALES dataset by AAclust approaches and *k-*free clustering models (**Fig. S1**). We first optimized *k*-optimized AAclust settings for seven models, such as hierarchical agglomerative clustering (HAC). The best performance was achieved by *k-*means when using merging, *min_cor*_*center*_, and *min_th*=0. Five of the seven *k*-optimized AAclust approaches outperformed *k-*free models (**Fig. 1d**), with the highest BIC scores around 25 clusters, declining linearly thereafter (**Fig. 1e**). Comparing all *k*-optimized against *k*-based approaches (with equal number of clusters) showed a significant (*P*<0.001) impact of merging on the clustering quality, improving BIC but worsening CH, while *k*-based approaches generally enhanced SC and CH (**Fig. S1**).

### 3.2 Evaluation of scale selection

We complied 24 benchmark datasets (**Fig. S2**), including 6 for ‘sequence prediction’ and 18 for ‘residue prediction’, to assess AAclust scale selections (**Fig. S3**). These selections were used as features for machine learning models, with their average accuracy (‘mean accuracy’) on the benchmark datasets indicating the quality of the scale set.

#### 3.2.1 Evaluation of AAclust scale selection

We evaluated *k*-based AAclust scale selections by obtaining scale sets of size ranging between 2 and 585 scales for seven clustering models (**Fig. S3**). The top-performing approaches were ranked for each dataset separately, showing similar mean accuracy values. The best-ranked models had the fewest scales (median: 103, inter quantile range (IQR): 50– 167), and the model rank positively correlated with the number of scales (Spearman’
ss correlation=0.16, *P<*0.05). Some clustering models, such as HAC (single), performed weakly in clustering (**Fig. 1d, e**) but well in prediction (**Fig. 1f**), while others exhibited the opposite trend, such as *k-*means.

AAclust-based strategies (*k*-optimized and *k*-based) significantly improved mean accuracy over the three baseline scale sets (**Fig. 1g, Fig. S4**). Notably, *k*-based approaches had a tad better performance than *k*-optimized approaches, albeit with higher variability (**Table S4)**. For most datasets and scale set ranges, *k*-based AAclust approaches outperformed randomly sampled sets (**Fig. S3**). However, pc-based sets were superior for ranges with n<=20 scales.

#### 3.2.2 Correlation of clustering quality and prediction performance

For the *k*-based AAclust approaches, we examined correlations between clustering quality, the number of scales, and the scale set quality (as quantified by machine learning model performance). The prediction performance was averaged across seven clustering models for each scale set size and benchmark dataset (MEAN_ACC_*dataset*). We then hierarchically clustered these 24 benchmark datasets into two groups (D1 and D2, **Fig. S5**).

We aggregated the model performance for D1 and D2 (‘ACC|D1’ and ‘ACC|D2’) and explored Pearson correlations across four scale set ranges. Correlations between the model performance and the number of scales were mainly positive, particularly for the 2–29 range and D2. For larger scale ranges, these correlations diverged, varying by clustering model and dataset. After min-max normalization, most clustering models achieved a normalized accuracy >=95% for D1 with few scales (e.g., 3 for *k-*means), while more than 35 were required for D2 (**Fig. 1h**). Remarkably, pc-based sets achieved optimal results with just 5 scales (*i*.*e*., the first 5 PCs) for D1, but required all 20 for D2 (**Fig. S6**).

#### 3.2.3 Effect of AAclust settings on prediction performance

We evaluated the impact of *k*-optimized AAclust settings on prediction performance for dataset groups D1 (ACC|D1) and D2 (ACC|D2). Generally, prediction performance was lower for D1 than for D2, and the best results for D2 were obtained with *min_th* between 0 and 0.6 without merging, where HAC (average) for D1 and *k-*means for D2 performed best (**Fig. S7**).

Analyzing cluster merging showed that *k*-based approaches performed significantly (*P*<0.001) lower for D1, while for D2, *k*-based and *k*-optimized (without merging) approaches were significantly better (*P<*0.001) than those using merging. Disabling the ‘Center’ parameter significantly (*P<*0.01–0.001) improved D2 performance. Assessing merging impact on min-max normalized accuracy values showed that smaller scale sets are preferred for D1 and larger for D2, consistent with prior results (**Fig. 1h**). Overall, our results emphasize that the optimal scale selection depends on the clustering model and the protein dataset.

### 3.3 AaclustTop60

We compiled the 60 best scale sets from all AAclust approaches into ‘AAclustTop60’ (**Fig. S8, Supplementary Table 3**). This collection comprises 48 top-ranked sets for 24 benchmark datasets (**Table S4**) and 12 top-ranked sets for dataset groups D1 and D2. We ranked these sets by average prediction performance and clustering quality, showing an anti-correlation (Pearson’
ss r=-0.77, P<0.01), whereby the number of scales correlated negatively with the former and positively with the latter. On average, sets in AAclustTop60 contained 125±121 scales (median: 98, IQR: 48–154) obtained using various AAclust settings. The variation in AAclustTop60 underlines that the optimal scale set depends on the protein prediction tasks.

## 4. Implementation

AAclust is integrated in AAanalysis, a Python framework for interpretable sequence-based protein prediction. Besides AAclust, AAanalysis will also provide the complete scale sets (SCALES), the 12 protein datasets, and the AAclustTop60.

For systematic optimization of sequence-based feature engineering using AAclust, we recommend the following steps:

1. Test all sets of AAclustTop60 as distinct features to establish baseline models and identify the best *k*-optimized AAclust settings, clustering models, and scale set ranges.
2. For the best clustering models, test (a) *k*-optimized AAclust approaches encompassing the best settings, and (b) *k*-based AAclust approaches within the optimal scale set range.
3. If the optimal range is within 2–20 scales, test pc-based scale sets.

Alternatively, AAclustTop60 scale sets could serve as initial population for genetic algorithms to optimize feature engineering [31]. Clustering models compatible with AAclust require a pre-defined number of clusters and should be implemented in scikit-learn or work accordingly.

## 5. Conclusion

We introduced AAclust, a clustering wrapper framework for selecting redundancy-reduced amino acid scale sets. Using Pearson correlation, AAclust optimizes the number of clusters and selects one scale per cluster. Our benchmarking experiments show that (a) no single ‘gold standard’ scale set exists, (b) the scale set size is a crucial and dataset-dependent optimization factor, and (c) AAclust scale selections significantly improve the performance of machine learning methods. Additionally, we collated the 60 best-performing scale sets (AAclustTop60) and provided a three-step application guide.

Although scale-based machine learning approaches have limitations [32], particularly in performance compared to deep learning-based protein embeddings (*i*.*e*., scale-like residue representation such as ProtT5 [33]), their advantage lies in their interpretability, which remains challenging for deep learning models [34]. Overall, AAclust tailors scale sets to specific protein prediction tasks, enabling systematic and interpretable sequence-based feature engineering.

## Supplementary Material

## SUPPLEMENTAL METHODS

### 1. Data representation and dataset collation

#### 1.1 General notation

We represent numerical data as *n*-dimensional vectors/arrays or *m* × *n* matrices containing *m* rows and *n* columns. Vectors are denoted by bold, lowercase letters, with individual elements indicated by indices, such as x_1_ referring to the first element of vector **x** = [x_1_, …, x_n_]. Matrices are represented by italic, bold, uppercase letters, with elements denoted by double indices, such as x_1,1_ referring to the first row, first column element in matrix ***X***:

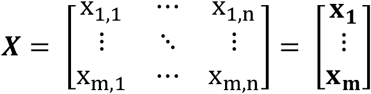

Sets of vectors and sets of sets or matrices are denoted by uppercase and bold uppercase letters, respectively. For example, a set of vectors C_1_ = {**x, y, z**} may be part of a larger set **C** = {C_1_, C_2_, …, C_n_}. Mathematical functions are indicated by lowercase italic letters, with parentheses enclosing their arguments, such as *min*(**x**) for computing the minimum value of **x**.

### 1.2 Dataset of amino acid scales

We obtained 566 physicochemical property scales for amino acids from the AAindex database (version 9.2) [1]. A further 86 scales related to the solvent accessible surface area (ASA) [2] and hydrophobicity of amino acids were manually collated from the literature (72 from Lins et al. [2] and 14 from Koehler et al. [3]) because of their general relevance for protein folding [4] and backbone dynamics [5]. After discarding scales due to missing values or complete redundancy, a set of 586 scales remained, which is referred to as SCALES (553 from AAindex, 21 from Lins et al., and 12 from Koehler et al.). For Lins et al. and Koehler at al., we created new scale ids by adopting the AAindex naming convention: first author’
ss last name, publication year, and the order of appearance in the publication (LINS030101, …, LINS030121 and KOEH090101, …, KOEH090112). Each scale was min-max normalized to the range of [0,1] (**Supplementary Table 1**).

### 1.3 Representation of amino acid scales

Property scales are represented as arrays **x** containing 20 numerical values, each corresponding to one of the 20 canonical amino acids: **x** = [x_1_, x_2_, … x_20_]. *m* Property scales are represented as an *m* × 20 feature matrix ***X***:

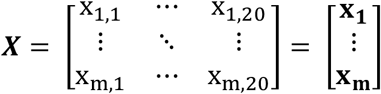

### 1.4 Scale sets for benchmarking

The scale sets selected by AAclust were compared against the following three groups of baseline scale sets:

- **standard**: Three ‘gold standard’ sets, two from previous studies comprising 7 [6,7] and 12 [8] scales (**Supplementary Table 1**), and all 586 scales from SCALES.
- **pc-based**: All scales from SCALES were transformed using principal component (PC) analysis into 20 principal components, each serving as a single scale, named P1 to P20.
- **random:** Subsets of varying size were assembled by randomly sampling scales from SCALES, a process repeated 10 times for each set size, to create random reference sets.

### 1.5 Datasets of protein sequences

We collated 12 protein sequence datasets, each previously utilized for binary classification problems (**Fig. S2**). 6 datasets were used to predict biological characteristics of entire protein sequences (e.g., protein solubility), while the other 6 were utilized for predicting properties of individual residues (e.g., solvent accessibility). We refer to these two groups as ‘sequence prediction’ and ‘residue prediction’ datasets, respectively. Each dataset was named based on its prediction task, either ‘SEQ_*dataset_name*’ or ‘AA_*dataset_name*’. **Table S1** offers an overview, with **Supplementary Table 2** providing additional details of these datasets.

For residue prediction, we considered contiguous amino acid segments of length *n*, centred on the target residue’s sequence position. Since the optimal window size is crucial for accurate predictions [9], we chose three window sizes (*n*=5, 9, 13), resulting in 18 residue prediction datasets (‘AA*n*_*dataset_name’*) and a total of 24 benchmark datasets. To enhance training efficiency and maintain comparability, each benchmark dataset included 400 randomly selected samples per class (*i*.*e*., proteins or residues), creating balanced datasets with 800 samples each.

**Table S1:**
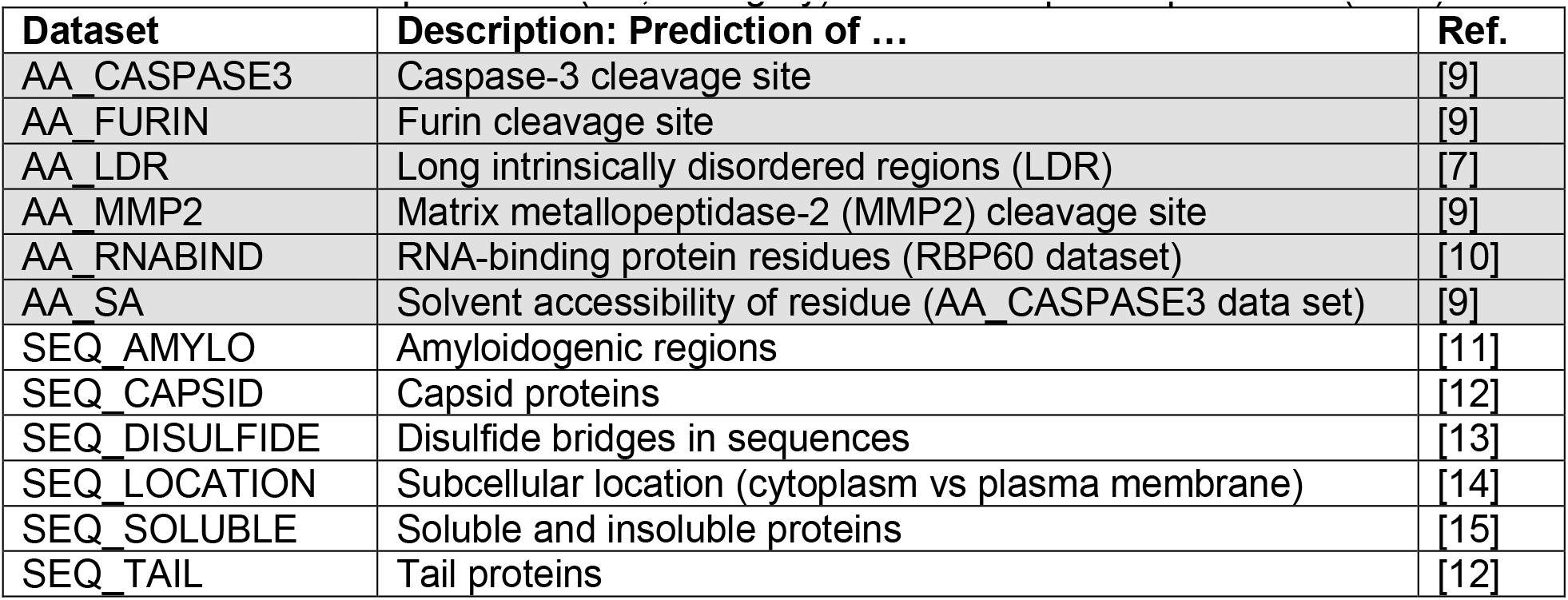
Overview of the protein datasets used for benchmarking of the AAclust scale selection at the residue prediction (AA; dark gray) or at the sequence prediction (SEQ) level.

| Dataset | Description: Prediction of ... | Ref. |
| --- | --- | --- |
| AA_CASPASE3 | Caspase-3 cleavage site | [9] |
| AA_FURIN | Furin cleavage site | [9] |
| AA_LDR | Long intrinsically disordered regions (LDR) | [7] |
| AA_MMP2 | Matrix metalloproteinase-2 (MMP2) cleavage site | [9] |
| AA_RNABIND | RNA-binding protein residues (RBP60 dataset) | [10] |
| AA_SA | Solvent accessibility of residue (AA_CASPASE3 data set) | [9] |
| SEQ_AMYLO | Amyloidogenic regions | [11] |
| SEQ_CAPSID | Capsid proteins | [12] |
| SEQ_DISULFIDE | Disulfide bridges in sequences | [13] |
| SEQ_LOCATION | Subcellular location (cytoplasm vs plasma membrane) | [14] |
| SEQ_SOLUBLE | Soluble and insoluble proteins | [15] |
| SEQ_TAIL | Tail proteins | [12] |

### 1.5 Representation of protein sequences

In the sequence prediction datasets, protein sequences were represented using *m* property scales. For each of the *m* scales, the corresponding amino acid values were assigned to each residue and averaged, resulting in an array with *m* average scale values. Consequently, a set of *n* protein sequences is described by an *n* × *m* matrix.

For the residue prediction datasets, a single residue was represented using a defined window size *n*, encompassing the central residue and its n-1 adjacent residues: (n-1)/2 at the N-terminal and (n-1)/2 at the C-terminal side. The residues within this window were represented by *m* property scales as before, generating an *n* × *m* matrix for *n* residues.

## AAclust: *k-*optimized clustering

### 2.1 Notation for clustering analysis

We will use the following notation to describe clustering measures and algorithms:

- **Observation (data point)**: Each observation is a vector with *n* real value attributes (or features), represented by a bold, lowercase letter, such as **x** = [x_1_, x_2_, …, x_n_]. In this work, observations refer to scales described by *n* = 20 numerical values corresponding to the 20 canonical amino acids. In general, AAclust could be used with any numerical vector.
- **Set of observations**: A set S of *m* observations is denoted as S = {**x**_1_, **x**_2_, …, **x**_m_}. They can be referred to by descriptive names such as for SCALES.
- **Cluster**: A cluster C is a set of *j* observations given as C = {**x**_1_, **x**_2_, …, **x**_j_} with C ⊆ S.
- **Set of clusters**: A set of *k* clusters is represented by bold letters as **C** = {C_1_, C_2,_ …, C_k_}, where each observation in S is assigned to exactly one cluster in **C**.
- **Number of clusters**: *k* is equal to the number of elements in **C** (*k* = |**C**|).
- ***Centroid***: A cluster center, denoted as *centroid*, is defined as the arithmetic mean of all observations within that cluster.
- ***Medoid***: A *medoid* is defined as the observation most similar to the *centroid* either based on Pearson correlation or distance measures such as Euclidean distance.
- **Clustering model**: A clustering model *CM* aims to partition S into *k* disjoint subsets, with each subset of S referred to as a cluster, and *k* being the number of clusters.

### 2.2 Idea of AAclust algorithm

AAclust is a versatile clustering wrapper [16,17] framework designed to work with clustering models that depend on a pre-defined number of clusters *k*. Rather than requiring *k* to be defined in advance, AAclust optimizes it automatically and selects one representative scale per cluster.

The input to AAclust consists of a clustering model and a set S of *m* observations, each with *n* attributes, represented as S = {**x**_1_, **x**_2_, …, **x**_m_} or an *m* × *n* feature matrix ***X***. While ***X*** could represent any feature matrix, the initial feature matrix for this study is a 586 × 20 matrix, comprising all 586 scales from SCALES. Ultimately, AAclust aims to find optimal subsets of S based on Pearson correlation.

### 2.3 Clustering models used in conjunction with the AAclust framework

AAclust works in conjunction with clustering models that use a pre-defined *k*, such as *k-*means [18] and hierarchical agglomerative clustering (HAC) [19,20], but not with models optimizing *k* internally, referred to as ‘*k* free’ models, such as DBSCAN [21] or OPTICS [22]. Here, the term ‘clustering model’ refers to AAclust-compatible models, unless stated otherwise. Alongside four variations of HAC, each using a different linkage method and considered as separate models, a total of seven models were tested with AAclust (see **Table S2**).

**Table S2:** Overview of the clustering models tested in conjunction with AAclust (dark grey) and *k-*free clustering models used for benchmarking.

| Method name | Short description | Ref. |
| --- | --- | --- |
| $k$-means | Partitional clustering based on within-cluster distance | [18] |
| Hierarchical agglomerative clustering (HAC) | Hierarchical clustering with four different types of linkage methods (average, complete, single, ward) | [19, 20] |
| Mini-batch $k$-means | Variation of $k$-means using mini-batches | [23] |
| Spectral clustering (Spectral) | Clustering with low-dimension graph embeddings for data grouping and structure identification | [24] |
| DBSCAN | Density clustering via distance between nearest points | [21] |
| OPTICS | Density clustering via distance between nearest points | [22] |
| BIRCH | Clustering feature tree on pairwise Euclidean distance | [25] |
| Affinity propagation | Clustering by passing messages between data points | [26] |

### 2.4 AAclust objective function based on two measures of minimum correlation

The objective function of AAclust relies on two distinct quality measures: *min_cor*_*all*_ and *min_cor*_*center*_. Both measures quantify the similarity between two observations/scales by employing the Pearson correlation coefficient. Given two vectors **x, y** with *n* pairs of sample points {(x_1_, y_1_), …, (x_n_, y_n_)}, the Pearson correlation *cor*(**x, y**) is defined as:

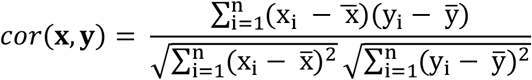

where the sample means are given by 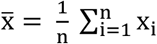 and 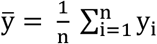.

Given a cluster C of *j* observations, represented as a feature matrix ***X***, the pairwise Pearson correlation matrix ***P*** is a symmetrical *j* × *j* matrix computed by *pairwise_cor(****X***):

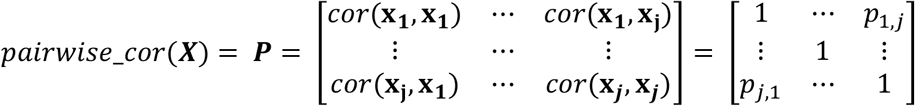

where *p*_*j1,j2*_ is the pairwise Pearson correlation between two observations ***j***_***1***_ and ***j***_***2***_, with real number value between -1 and 1.

The first quality measure *min_cor*_*all*_(***X***) is defined as the minimum of the pairwise Pearson correlation matrix ***P***, computed as:

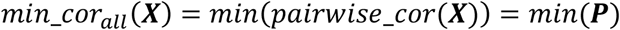

To define the second quality measure, we first define a cluster center as the vector of mean values for each attribute (*centroid*), calculated over all observations in a cluster [27]. The *centroid* can be computed for a feature matrix ***X*** with *m* observations and *n* attributes as:

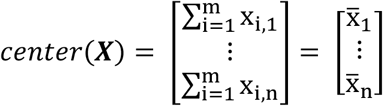

The Pearson correlation between an observation **x** from a feature matrix ***X*** and the center of ***X*** is given by:

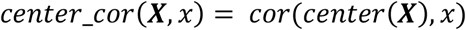

The second quality measure *min_cor*_*center*_(***X***) for a feature matrix ***X*** with *m* observations is defined as the minimum of the Pearson correlation with the center, computed as:

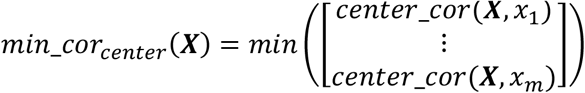

Both *min_cor*_*all*_ and *min_cor*_*center*_ can be used by AAclust to estimate and optimize *k*. For a set of *k* clusters in **C** = {C_1_, …, C_k_}, the objective function *min_cor*(**C**) is computed as:

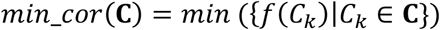

where *f* can be *min_cor*_*all*_ or *min_cor*_*center*_. The objective function of AAclust computes the minimum of the Pearson correlation for each cluster, using as quality measure either the minimum within-cluster correlation (*min_cor*_*all*_) or the minimum correlation with the cluster center (*min_cor*_*center*_), referred to as minimum ‘with-center’ correlation. Considering a clustering model *CM*, we can give the objective function by *min_cor*(***X***, *k, CM*), where ***X*** is a feature matrix and *k* the number of clusters.

AAclust aims to find the smallest number of clusters such that the minimum correlation for each cluster is greater than or equal to a defined minimum correlation threshold *min_th*. For example, if *min_th* is set to 0.3 and the optimization is performed using *min_cor*_*all*_, the minimum within-cluster Pearson correlation for each cluster will be at least 0.3, meaning that all observations assigned to the same cluster correlate at least weakly (as shown in **Fig. 1a**).

### 2.5 Algorithm of AAclust

AAclust is a wrapper for a clustering model *CM* that uses a pre-defined *k*. It aims to partition a given feature matrix ***X*** of *m* observations/scales into a set of *k* clusters **C**. AAclust estimates and optimizes *k* by the objective function *min_cor(***C**) (or *min_cor*(**X**, *k*, CM)) with either *min_cor*_*all*_ or *min_cor*_*center*_ employed as a quality measure.

As shown in **Fig. 1a** and outlined in **Algorithm 1–4**, the AAclust algorithm comprises the following three steps:

#### 1. *Estimation of k* (Algorithm 1)

The lower bound of *k* is estimated by computing *min_cor*(**C**) in 10% increments of the total number of observations. The process begins with *k*=1 (lines 1–7) and continues until *min_cor*(**C**) exceeds the user-defined threshold *min_th* (lines 7– 11), which represents the minimum acceptable within-cluster Pearson correlation over all clusters. The second highest lower bound of *k* is selected (lines 12–13) to provide a conservative estimation of the true lower bound. For example, given *min_th*=0.3 and a set of 100 scales, *min_cor*(**C**) would be computed for *k* = 1, 10, 20, 30 …, 100 as long as *min_cor*(**C**) is less than 0.3. This would result in a vector of results **c** such as **c**=[-0.5, 0.1, 0.2, 0.3]. In this case, the second highest lower bound of *k* would be 10 (with *min_cor*(**C**)=0.1). This procedure has been empirically optimized to find a good trade-off between efficiency and correctness.

#### 2. *Recursive clustering* (Algorithm 2)

The number of clusters *k* is incrementally optimized using a recursive clustering procedure. For *m* observations, an initial step size *s* is chosen between 1 to 5 by using *s* = max(1, min(*floor*(*m*/10),5)), where *floor*(*x*) returns the greatest natural value less than or equal to *x* (lines 1–2). The estimated value of *k* is then incremented stepwise by adding s (*k* = *k + s*), and the minimum within-cluster Pearson correlation (*min_cor*(**C**)) is computed for the updated value of *k* (lines 4–5). If the initial step size is not 1, this process is repeated until *min_cor*(**C**)>=*min_th* (line 6). Once *min_cor*(**C**) exceeds *min_th*, the value of *k* is decreased by *k* = max (1, *k − s ×* 2), *s* is set to 1, and *min_cor*(**C**) is calculated for the decreased value of *k* (lines 7–9). This adjustment of *k* is performed to ensure that the true optimum of *min_cor*(**C**) is found in the subsequent iterations. Finally, *k* is incremented stepwise by 1 until either *min_cor*(**C**) exceeds *min_th* or each observation forms a unique cluster (*k*=*m*; lines 3–5). This procedure was empirically optimized.

#### 3. *Cluster merging* (Algorithm 3)

Optionally, smaller clusters can be merged into bigger clusters, meaning that all observations of the small cluster are reassigned to bigger clusters. Small clusters are defined as clusters with *n_max<*=5 observations. The process begins with *n_max*=1 and continues until *n_max*=5, where each small cluster is merged into each bigger cluster, and the minimum within-cluster Pearson correlation (*min_cor*(**C**)) is computed for each merged cluster combination (lines 2–8). If *min*_cor(**C**) exceeds *min_th*, the maximum within-cluster Euclidean distance *d_max*(**C**) for the combined cluster is computed as a quality measure for merging (lines 9–11). Smaller clusters are then merged into the bigger cluster with the lowest *d_max*(**C**) (lines 12–13). Alternatively, *min*_cor(**C**) can be used instead of *d_max*(**C**).

In summary, AAclust (**Algorithm 4**) optimizes *k* through estimation of its lower bound, recursive clustering, and optional merging of small clusters into bigger ones. The entire process is guided by the goal of maximizing the overall within-cluster or with-center Pearson correlation to meet a user-defined minimum correlation threshold (*min_th*).

##### Algorithm 1

Estimate*k*

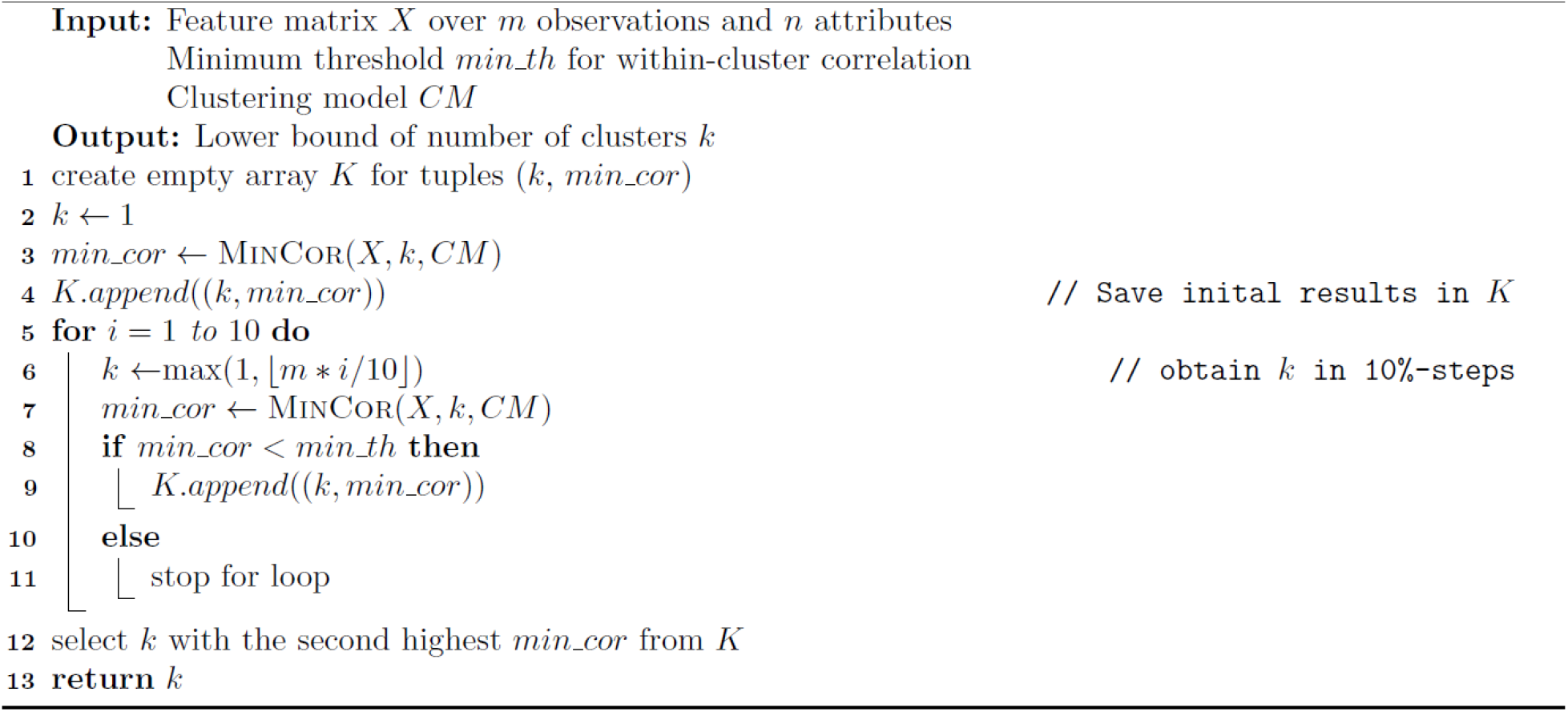

##### Algorithm 2

Optimize*k*

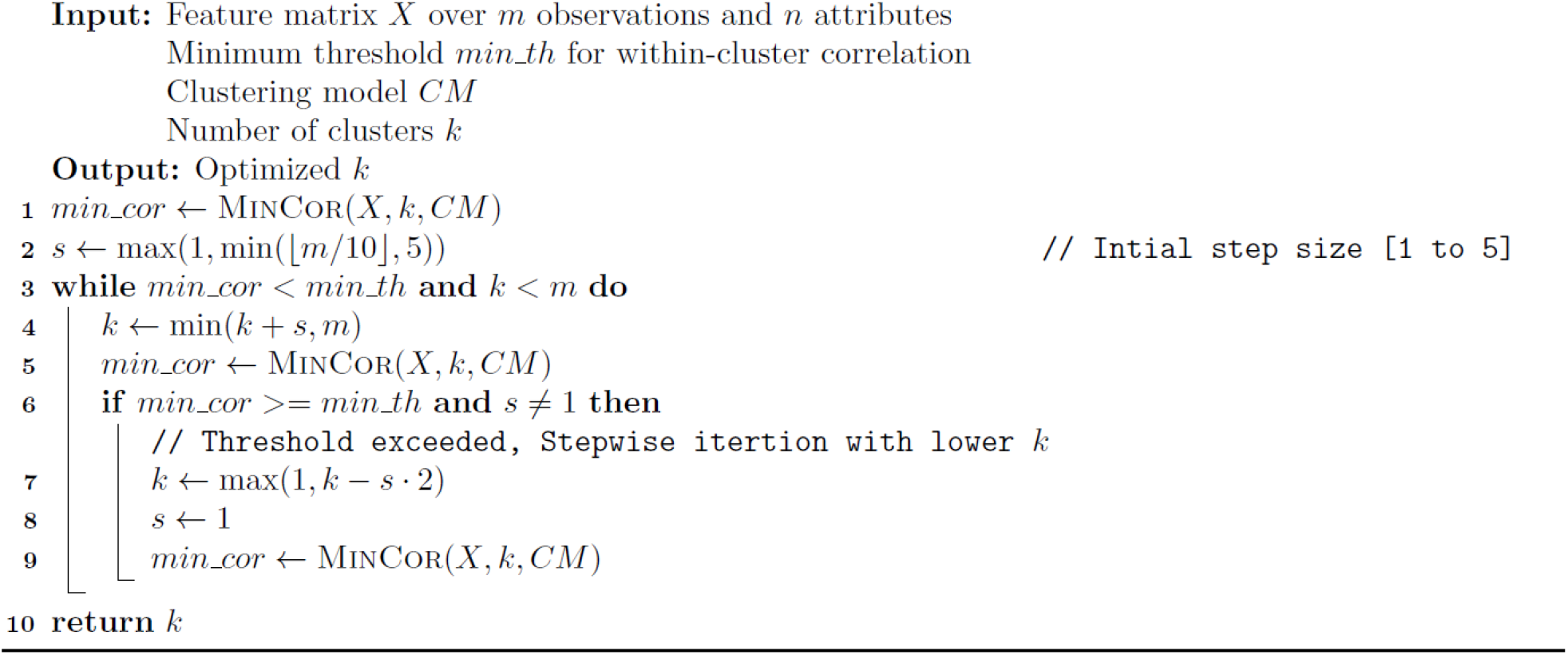

##### Algorithm 3

MergeGlusters

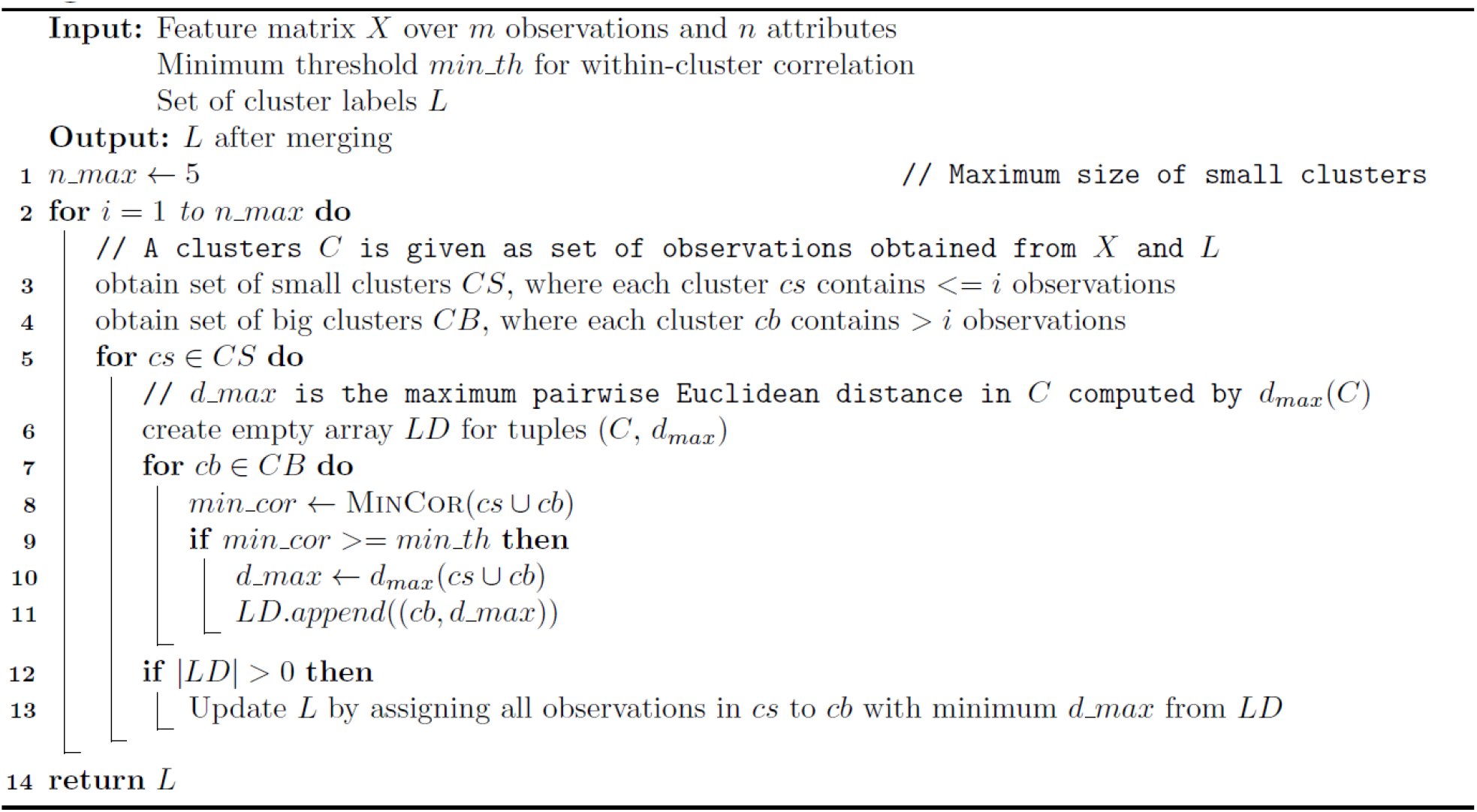

##### Algorithm 4

AAclust

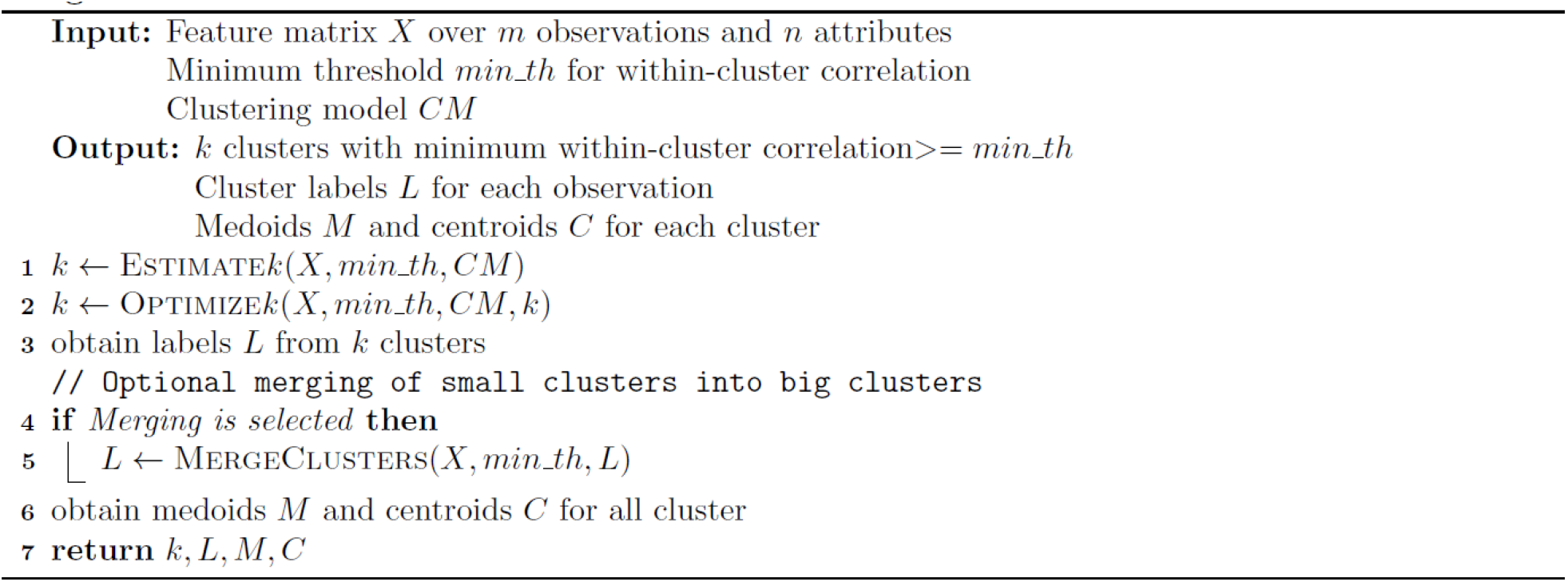

### 2.7 AAclust for scale selection

AAclust can identify redundancy-reduced subsets of scales. Similar to other bioinformatics tools such as REVIGO [28] and CD-Hit [29], AAclust first performs an optimized clustering procedure. It then selects for each cluster a single representative scale (*medoid*) closest to its *centroid*, using Pearson correlation or Euclidean distance. Users can determine the number of clusters *k* (and thus selected scales) in two ways (**Fig. 1b**):

- ***k-*optimized**: *k* is automatically optimized by AAclust as described before.
- ***k-*based**: *k* is directly specified by the user.

When comparing both approaches, the *k* value optimized in the *k-*optimized approach was used for *k-*based approaches with the same clustering model, such that their number of clusters matched.

### 2.8 AAclust parameters

AAclust is controlled by is three parameters (**Fig. 1a**):

- ***min_th***: Sets the minimum Pearson correlation threshold (0 to 1, default 0.3).
- **Center**: Determines whether *min_th* applies to the cluster center (true) or all members (false), using either *min_cor*_*center*_ or *min_cor*_*all*_, respectively.
- **Merge**: Enables (true) or disables (false) the optional merging step, utilizing Pearson correlation or distance measures such as Euclidean distance.

We recommend using the phrase ‘minimum within-cluster/with-center Pearson correlation of X’ to describe the usage of *min_th*=X with *min_cor*_*center*_ or *min_cor*_*all*_, respectively.

We tested 60 *k-*optimized AAclust approaches, varying *min_th* (0, 0.1, …, 0.9), merging options (no merging, merging with Euclidean distance or Pearson correlation), and the objective function. For clarity, our results (**Fig. S1c, Fig. S7a**) highlight only certain thresholds (*min_th*=0, 0.3, 0.6, 0.9) and Euclidean-based merging due to its slightly superior performance.

## 3. Quality measures used for the evaluation of clustering and scale selection

### 3.1 Quality measures for clustering

The clustering performance of AAclust was evaluated using established quality measures [30–32], including the silhouette coefficient (SC) [33,34], the Calinski Harabasz score (CH) [35], and the Bayesian information criterion (BIC) [36]. While the BIC strives to find the simplest data-explaining model, the SC and CH assess two primary clustering aspects (**Fig. S1a**) [37]:

- **Cohesion/compactness**: This reflects how closely related or similar elements within the same cluster are. Cohesion can be measured through either the within-cluster/intra-cluster distance (*i*.*e*., the average of all pairwise distances inside a cluster) or the within-cluster dispersion (*i*.*e*., the sum of squared distances from data points to the *centroids*).
- **Separation**: This quantifies how distinct or scattered apart clusters are from each other. Separation can be measured using metrics such as the between-cluster/inter-cluster distance (*i*.*e*., the distances between the *centroids* of each pair of clusters), the nearest-cluster distance (*i*.*e*., the average distance of a data point with all data points from the closest cluster), or the between-cluster dispersion (*i*.*e*., the sum of squared distances between the *centroids* of all clusters and the overall data mean).

Clustering quality measures were obtained using sklearn.metrics.silhouette_score, sklearn.metrics.calinski_harabasz_score, and a self-devised BIC implementation adopted from [38]. This BIC version was modified to align with the SC and CH, such that higher values signify better clustering, contrary to the conventional BIC interpretation favoring lower values. Further details can be found in **Table S3**, adapted from [32].

**Table S3.** Quality measures used for evaluation of clustering algorithms.

| Name | Best if ... | Range | Abbreviation | Reference |
| --- | --- | --- | --- | --- |
| Silhouette coefficient | High | [-1, 1] | SC | [33,34] |
| Calinski Harabasz score | High | [0, ∞] | CH | [35] |
| Bayesian information criterion | High | [-∞, ∞] | BIC | [36] |

#### Computation of the silhouette coefficient

The SC is a measure of how well data points are grouped within clusters based on silhouette values [33]. The silhouette value [34] of an observation quantifies its similarity to the other objects in its own cluster (*cohesion*) compared to objects in the closest other cluster (*separation*). The cohesion and separation of a cluster are calculated by the mean within-cluster distance and the mean nearest-cluster distance [34], respectively. The silhouette value for an observation **x**, which belongs to cluster A but not to cluster B [34], is defined as:

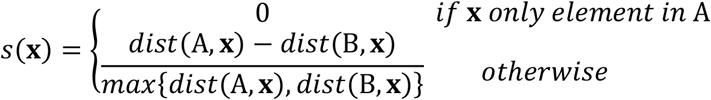

where *dist*(A, **x**) is the mean within-cluster distance of **x**, calculated as the average distance between **x** and all other points in A (excluding **x**), using a distance metric such as Euclidean distance:

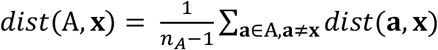

and *dist*(B, **x**) is the mean nearest-cluster distance, calculated as the average distance between **x** and all data points in the cluster B (with the smallest mean distance to **x** among all clusters in set **C**):

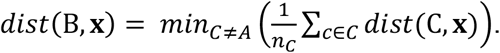

The *SC* is then calculated as the average of all silhouette values for *n* data points across a set of clusters ***C***:

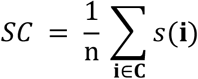

#### Computation of the Calinski Harabasz score

The CH is a metric to evaluate the quality of a clustering solution. It is calculated as the ratio between the between-cluster dispersion and the within-cluster dispersion, where dispersion is defined as the sum of distances squared. Given a set *n* data points S that has been partitioned into *k* clusters, the *CH* is defined [35] as:

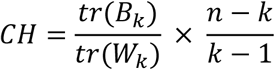

where *tr(B*_*k*_) is the trace of the between-cluster dispersion matrix and *tr(W*_*k*_) is the trace of the within-cluster dispersion matrix, defined as:

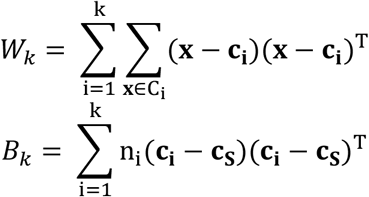

where C_*i*_ is a set with n_*i*_ data points or the *i*-th cluster of S (C_*i*_ ⊂ S), **c**_**i**_ is the centroid of cluster C_*i*_, and **c**_**S**_ is the center of S.

#### Computation of the Bayesian information criterion

The BIC is used to determine the simplest model that explains the data. It tries to minimize the within-cluster distortion, which is calculated as the average squared distance between all points and their centroid. Given a set S of *n* data points, each with *m* features, that has been partitioned into *k* clusters, *BIC* is defined as follows [38]:

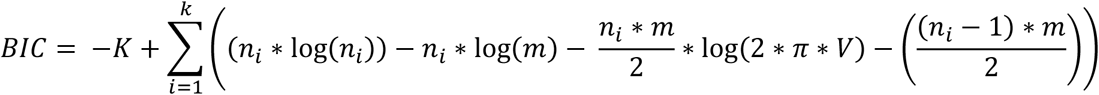

where n_*i*_ is the number of data points in cluster C_*i*_, with C_*i*_ ⊂ S. The constant *K* and the variance over all clusters *V* are defined as:

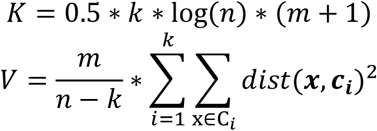

where *dist*(**a, b**) is the Euclidean distance between data points **a** and **b**, and **c**_***i***_ is the centroid of cluster C_*i*_. For a detailed derivation and definition of the BIC, see [38,39].

### 3.2 Machine learning models

To evaluate the performance of the scale sets, three machine learning classification models with default settings were utilized: random forest, support vector machine, and logistic regression classifier as implemented in sklearn.ensemble.RandomForestClassifier, sklearn.svm.SVC, and sklearn.liner_model.LogisticRegression, respectively. To prevent model-dependent bias, we averaged the performance of the three models.

### 3.3 Evaluation procedure for scale selections

The performance of the scales selected by AAclust were evaluated (**Fig. S3a**) by comparing them with ‘standard’, ‘pc-based’, and ‘random’ scale sets (see ‘1.4 Scale sets for benchmarking’). These served as features for training machine learning models in a binary classification approach on benchmark datasets. Model performance was assessed using accuracy (ACC), defined as: ACC= (TP *+* TN)/(TP *+* TN *+* FP *+* FN), where TP, TN, FP, FN represent the number of true positives, true negatives, false positives, and false negatives, respectively. Additional metrics were omitted due to the balanced nature of our benchmark datasets (see ‘1.5 Datasets of protein sequences’). A five-fold cross validation was conducted and the average accuracy (‘mean accuracy’) over all folds and three models was used as the quality measure for scale sets.

### 3.4 Statistical testing

To perform parametric testing of multiple groups, we utilized the one-way ANOVA with Sidak post hoc test as implemented in statsmodels.stats.multicomp.Multicomparison. The normality assumption was tested using the Shapiro-Wilk test as implemented in scipy.stats.shapiro. For independent t-tests, the scipy.stats.ttest_ind implementation was used. To perform nonparametric testing of paired data, the paired Wilcoxon signed-rank test was used, which was implemented in scipy.stats.wilcoxon. Pearson correlation was computed using the scipy.stats.pearsonr implementation.

## SUPPLEMENTAL RESULTS

### 4. Evaluation of clustering quality

To assess AAclust clustering quality, we clustered the 586 property scales from SCALES and evaluated the results using SC, CH, and BIC (**Fig. S1a, 1c**). The clustering was performed using *k-*free models and AAclust (*k-*optimized and *k-*based) with seven models, including *k-*means, MiniBatch *k-*means, Spectral clustering and HAC with four linkage methods.

We first assessed 60 AAclust parameter combinations (*k-*optimized), varying *min_th*, the merging step (example in **Fig. S1b**), and the objective function (see ‘2.8 AAclust parameters’). The best results were achieved using *k-*means (**Fig. S1c**, red squares) with *min_th*=0, *min_cor*_*center*_ (Center=True), and merging (Merge=True). Merging led to evenly sized clusters, with Euclidean distance outperforming Pearson correlation slightly (**Supplementary Table 3**). Next, we compared the best *k*-optimized AAclust models to three *k-*free models, including Birch, DBSCAN, and OPTICS (**Fig. 1d**). The best results were obtained by AAclust with *k-*means and HAC (ward). Density-based clustering models (DBSCAN and OPTICS) resulted in unclustered scales, which were reassigned to one cluster to maintain comparability. The impact of merging was evaluated by comparing the *k-*optimized results for the best models (with and without merging) against *k-*based approaches (**Fig. S1d**, best model (*k*-means) in red). Based on the results of the 60 AAclust parameter combinations, merging significantly improved BIC but worsened SC (*P<*0.001, one-way ANOVA, Sidak post hoc test), while *k-*based approaches generally improved SC and CH, though not significantly (**Fig. S1e**).

Evaluating the seven models using the *k-*based AAclust approach across 2–200 clusters revealed optimal BIC scores around 25 clusters, declining thereafter. In contrast, SC scores remained constantly high, while CH scores peaked at a low cluster number and declining starting from 2 clusters (**Fig. 1e**). An outlier was HAC (single)—likely due to its preference for single-datapoint clusters, leading to noisy results at low cluster numbers—which performed worse in all quality measures but showed rising BIC and SC scores after 100 clusters.

### 5 Evaluation of scale selection

#### 5.1 General evaluation workflow

AAclust reduces scale set redundancy by clustering and selecting one representative scale per cluster, resulting in *k* selected scales (**Fig. 1a, b**). These scale sets served as features for machine learning models (see ‘3.2 Machine learning models’) and were evaluated on accuracy (ACC) across 24 benchmark datasets (**Fig. S2**). In our testing workflow (**Fig. 1c**), we evaluated the prediction performance of the machine learning models using AAclust scale selections against other baseline scale sets, including ‘gold standard’, pc-based, and randomly sampled sets (see ‘1.4 Scale sets for benchmarking’). To assess the correlation between the prediction performance and the clustering quality, we hierarchically clustered all datasets, resulting in two groups (D1, D2), and aggregated the prediction performance for each group separately (ACC|D1, ACC|D2).

#### 5.2 Evaluation of AAclust scale selection

To evaluate AAclust scale selections (**Fig. S3a**), we employed 7 clustering models with the *k-*based AAclust approach on ranges comprising 2–585 scales, resulting in 4088 scale sets. These served as features for machine learning models across 24 benchmark datasets, using the mean accuracy as performance measure of the scale sets and clustering approaches.

We analyzed the prediction performance within specific scale set ranges and benchmark datasets to identify and rank the best-performing scale sets. The performance was comparable within each dataset (**Fig. S3b**). The highest-ranked models had the fewest number of scales (median: 103, inter quantile range (IQR): 50–167), and the model rank positively correlated (Spearman’
ss correlation=0,16, *P<*0.05) with the number of scales (**Fig. S3c**).Some clustering models, such as HAC (single), performed weakly in clustering (**Fig. 1d, e**) but well in prediction (**Fig. 1f**), while others exhibited the opposite trend, such as *k-*means and HAC (ward).

For most of the 24 benchmark datasets, *k-*based scale sets significantly outperformed randomly sampled scales for ranges of 2–60 and 2–585 scales (**Fig. S3d**), considering pairs of mean accuracy values for the same number of scales. However, pc-based scale sets surpassed AAclust-based scale sets for ranges of 2–20 scales in most datasets.

We then compared the top-ranked models with the best-performing models for the three baseline sets of ‘gold standard’, pc-based, and randomly selected scales. Considering pairs of mean accuracy values for the same benchmark dataset, AAclust approaches had a significantly better mean accuracy (*p*<0.1–0.001, paired Wilcoxon signed-rank test, Benjamini Hochberg correction [40]; **Fig. 1g, Fig. S4**). We assessed *k-*based against *k-*optimized AAclust approaches, both showing a similar mean accuracy (73.3±1.0% vs 73.3±0.6%), yet *k-*based approaches had a much higher variability (16.3% vs 3.5% max-min accuracy; see **Table S4**). Overall, AAclust-based scale sets outperformed baseline scale sets, except for ranges with n<=20 scales, where pc-based sets were superior.

**Table S4:**
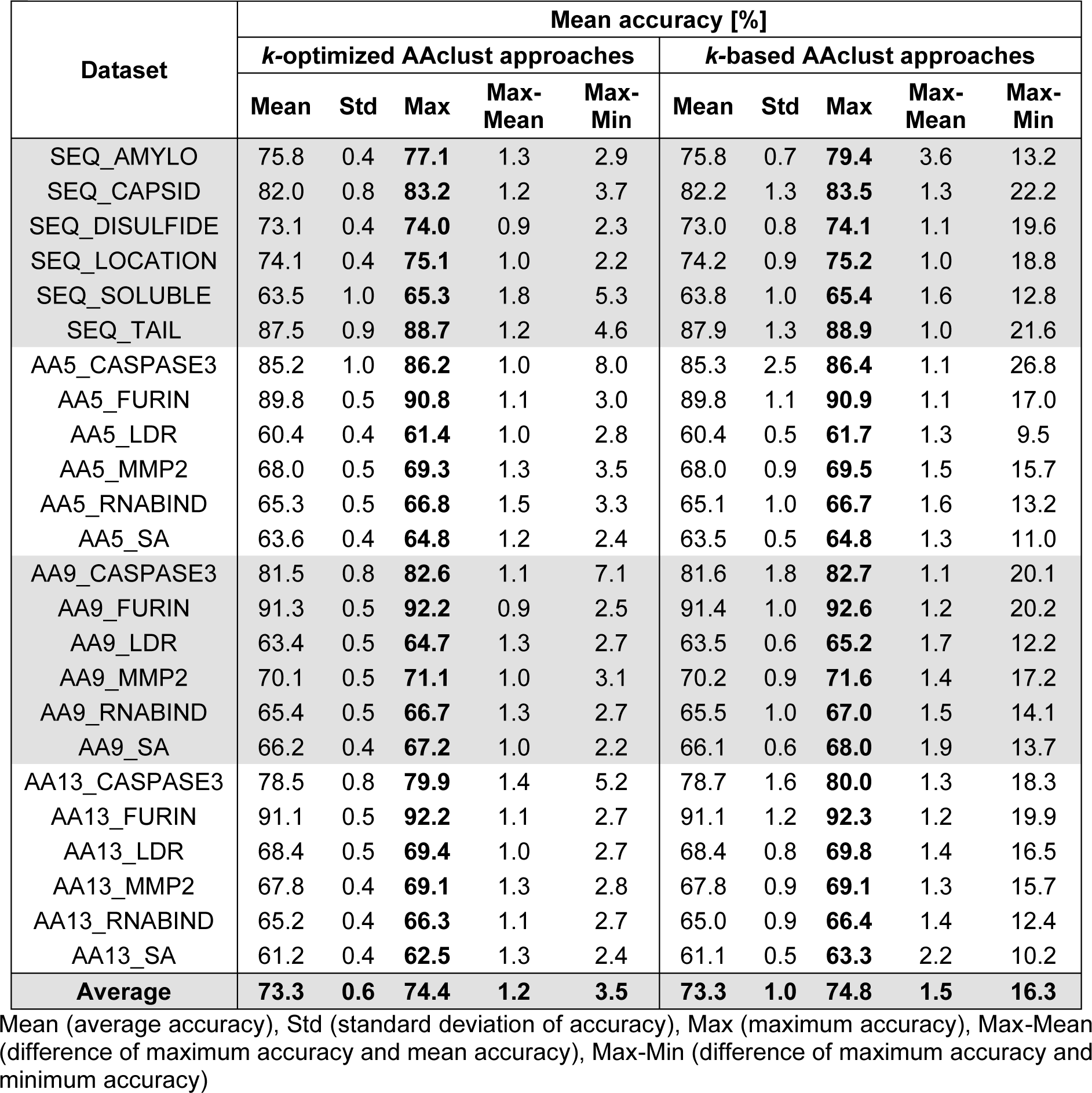
Mean accuracy for 24 benchmark datasets (6 at sequence prediction level (SEQ) and 18 at residue prediction level (AA) with window sizes 5, 9, 13), comparing *k-*optimized and *k-*based AAclust approaches. The results for the best-performing models are shown in the ‘Max’ column, indicated in bold. The average results are given in the last row.

| Dataset | Mean accuracy [%] |  |  |  |  |  |  |  |  |  |
| --- | --- | --- | --- | --- | --- | --- | --- | --- | --- | --- |
| | $k$-optimized AAclust approaches | | | | | $k$-based AAclust approaches | | | | |
|  | Mean | Std | Max | Max-Mean | Max-Min | Mean | Std | Max | Max-Mean | Max-Min |
| SEQ_AMYLO | 75.8 | 0.4 | **77.1** | 1.3 | 2.9 | 75.8 | 0.7 | **79.4** | 3.6 | 13.2 |
| SEQ_CAPSID | 82.0 | 0.8 | **83.2** | 1.2 | 3.7 | 82.2 | 1.3 | **83.5** | 1.3 | 22.2 |
| SEQ_DISULFIDE | 73.1 | 0.4 | **74.0** | 0.9 | 2.3 | 73.0 | 0.8 | **74.1** | 1.1 | 19.6 |
| SEQ_LOCATION | 74.1 | 0.4 | **75.1** | 1.0 | 2.2 | 74.2 | 0.9 | **75.2** | 1.0 | 18.8 |
| SEQ_SOLUBLE | 63.5 | 1.0 | **65.3** | 1.8 | 5.3 | 63.8 | 1.0 | **65.4** | 1.6 | 12.8 |
| SEQ_TAIL | 87.5 | 0.9 | **88.7** | 1.2 | 4.6 | 87.9 | 1.3 | **88.9** | 1.0 | 21.6 |
| AA5_CASPASE3 | 85.2 | 1.0 | **86.2** | 1.0 | 8.0 | 85.3 | 2.5 | **86.4** | 1.1 | 26.8 |
| AA5_FURIN | 89.8 | 0.5 | **90.8** | 1.1 | 3.0 | 89.8 | 1.1 | **90.9** | 1.1 | 17.0 |
| AA5_LDR | 60.4 | 0.4 | **61.4** | 1.0 | 2.8 | 60.4 | 0.5 | **61.7** | 1.3 | 9.5 |
| AA5_MMP2 | 68.0 | 0.5 | **69.3** | 1.3 | 3.5 | 68.0 | 0.9 | **69.5** | 1.5 | 15.7 |
| AA5_RNABIND | 65.3 | 0.5 | **66.8** | 1.5 | 3.3 | 65.1 | 1.0 | **66.7** | 1.6 | 13.2 |
| AA5_SA | 63.6 | 0.4 | **64.8** | 1.2 | 2.4 | 63.5 | 0.5 | **64.8** | 1.3 | 11.0 |
| AA9_CASPASE3 | 81.5 | 0.8 | **82.6** | 1.1 | 7.1 | 81.6 | 1.8 | **82.7** | 1.1 | 20.1 |
| AA9_FURIN | 91.3 | 0.5 | **92.2** | 0.9 | 2.5 | 91.4 | 1.0 | **92.6** | 1.2 | 20.2 |
| AA9_LDR | 63.4 | 0.5 | **64.7** | 1.3 | 2.7 | 63.5 | 0.6 | **65.2** | 1.7 | 12.2 |
| AA9_MMP2 | 70.1 | 0.5 | **71.1** | 1.0 | 3.1 | 70.2 | 0.9 | **71.6** | 1.4 | 17.2 |
| AA9_RNABIND | 65.4 | 0.5 | **66.7** | 1.3 | 2.7 | 65.5 | 1.0 | **67.0** | 1.5 | 14.1 |
| AA9_SA | 66.2 | 0.4 | **67.2** | 1.0 | 2.2 | 66.1 | 0.6 | **68.0** | 1.9 | 13.7 |
| AA13_CASPASE3 | 78.5 | 0.8 | **79.9** | 1.4 | 5.2 | 78.7 | 1.6 | **80.0** | 1.3 | 18.3 |
| AA13_FURIN | 91.1 | 0.5 | **92.2** | 1.1 | 2.7 | 91.1 | 1.2 | **92.3** | 1.2 | 19.9 |
| AA13_LDR | 68.4 | 0.5 | **69.4** | 1.0 | 2.7 | 68.4 | 0.8 | **69.8** | 1.4 | 16.5 |
| AA13_MMP2 | 67.8 | 0.4 | **69.1** | 1.3 | 2.8 | 67.8 | 0.9 | **69.1** | 1.3 | 15.7 |
| AA13_RNABIND | 65.2 | 0.4 | **66.3** | 1.1 | 2.7 | 65.0 | 0.9 | **66.4** | 1.4 | 12.4 |
| AA13_SA | 61.2 | 0.4 | **62.5** | 1.3 | 2.4 | 61.1 | 0.5 | **63.3** | 2.2 | 10.2 |
| **Average** | **73.3** | **0.6** | **74.4** | **1.2** | **3.5** | **73.3** | **1.0** | **74.8** | **1.5** | **16.3** |

Mean (average accuracy), Std (standard deviation of accuracy), Max (maximum accuracy), Max-Mean (difference of maximum accuracy and mean accuracy), Max-Min (difference of maximum accuracy and minimum accuracy)

#### 5.3 Correlation of clustering quality and prediction performance

We explored *k-*based AAclust approaches for correlations between clustering quality, scale set size, and scale set quality (as quantified by machine learning model performance). Performance was averaged across seven clustering models for each scale set size and benchmark dataset (‘MEAN_ACC_*dataset*’), and Pearson correlations were used to hierarchically cluster the 24 benchmark datasets (**Fig. S5a**) into two groups (D1 and D2; **Fig. S5b**). Clustering quality measures showed negative (SC, BIC) or no correlation (CH) with model performance, while the number of scales positively correlated, especially for D1.

We aggregated performances for D1 and D2 (‘ACC|D1’ and ‘ACC|D2’) for each clustering model, assessing correlations with the number of scales and BIC across four scale set ranges (2–29, 2–100, 2–200, 2–585; **Fig. S5c**). The 2–29 range showed positive correlations (stronger in D2), except for HAC (single). For larger ranges, correlations varied, with positive correlations for the number of scales and negative for BIC. An analysis of two representative datasets (SEQ_AMYLO and SEQ_CAPDSID) and two clustering models (*k-*means and HAC (average); **Fig. S5d**) illustrates that accuracy mainly positively correlates with the number of scales, while its correlation with quality measures varies by model and dataset.

Next, we examined the performance of clustering models by min-max normalizing their average accuracy for D1 and D2, determining the minimum number of scales needed for a normalized accuracy >=95% (**Fig. 1h**). Most models achieved this with few scales for D1 (e.g., 3 for *k-*means) but needed over 35 for D2. For D1, pc-based scales required only 5 scales (*i*.*e*., the first 5 PCs), but 20 scales were needed to achieve a performance >= 95% for D2 (**Fig. S6**).

In conclusion, no systematic relationship was found between clustering quality and scale set quality, but a positive correlation was observed between the model performance and the scale number, particularly in smaller scale set ranges, all model- and dataset-specific.

#### 5.4 Effect of AAclust settings on prediction performance

We evaluated *k-*optimized AAclust settings across dataset groups D1 and D2, testing 60 combinations per clustering model by varying *min_th*, merging options, and the objective function (see ‘2.8 AAclust parameters’). Performance was lower for D1, with no apparent correlation with settings, whereas the best results for D2 were achieved with *min_th* values between 0 and 0.6 without merging. Best performing models were *k-*means for D1 and HAC (average) for D2 (**Fig. S7a**, red squares).

Analyzing cluster merging, we found that *k-*based approaches performed significantly lower for D1 (*P<*0.001, one-way ANOVA, Sidak post hoc test), while for D2, *k-*based and *k-*optimized (without merging) approaches were significantly better (*P<*0.001) than those using merging (**Fig. S7b**). Using the *min_cor*_*all*_ objective function (‘Center’=False) significantly improved performance for D2 (*P*<0.01–0.001, independent t-test, Bonferroni correction; **Fig. S7c**).

Finally, we observed that merging increased the accuracy for D1 (57% to 63%) but decreased it for D2 (86% to 75%), where all merging-based models with fewer than 100 scales performed below 80% (**Fig. S7d**, best-performing models in red). This finding aligns with the result for the *k-*based AAclust approaches, indicating that smaller scale sets were preferred for D1 and larger scale sets for D2 (**Fig. 1h**). Overall, our results emphasize that the optimal scale selection depends on the clustering model and the protein dataset.

### 6. Creation and evaluation of AAclustTop60

We compiled the ‘AAclustTop60’, a selection of the 60 best scale sets from all *k-*optimized (n=420) and *k-*based (n=4088) AAclust approaches (**Fig. S8a**; **Supplementary Table 3, 4**). It encompasses 48 top-ranked scale sets based on the protein benchmark datasets and 12 sets derived from the aggregated prediction performance of groups D1 and D2 (**Fig. 1h**). Over 1/2 of the scale sets were obtained using HAC with single and complete linkage, followed by *k-*means (**Fig. S8b**), with only three property scales occurring in 2/3 of the 60 sets (**Fig. S8c**).

The scale sets were ranked based on their average prediction performance and clustering quality (BIC, CH, SC), both showing an anti-correlation (Pearson’
ss r=-0.77, *P*<0.01; **Fig. S8d**), while the clustering-based ranking positively correlated with the number of scales (Pearson’s r=0.3, *P<*0.001). AAclustTop60 sets contain on average 125±121 scales (median: 98, IQR: 48–154), with *k-*based approaches showing better clustering and prediction rankings (**Fig. S8e**). Comprising 36 *k-*based and 24 *k-*optimized approaches, AAclustTop60 shows variations for the used correlation thresholds and merging settings (**Fig. S8f**). These findings demonstrate the diversity within AAclustTop60, emphasizing that the optimal scale set must be tailored to the specific protein prediction tasks.

**Fig. S1.**
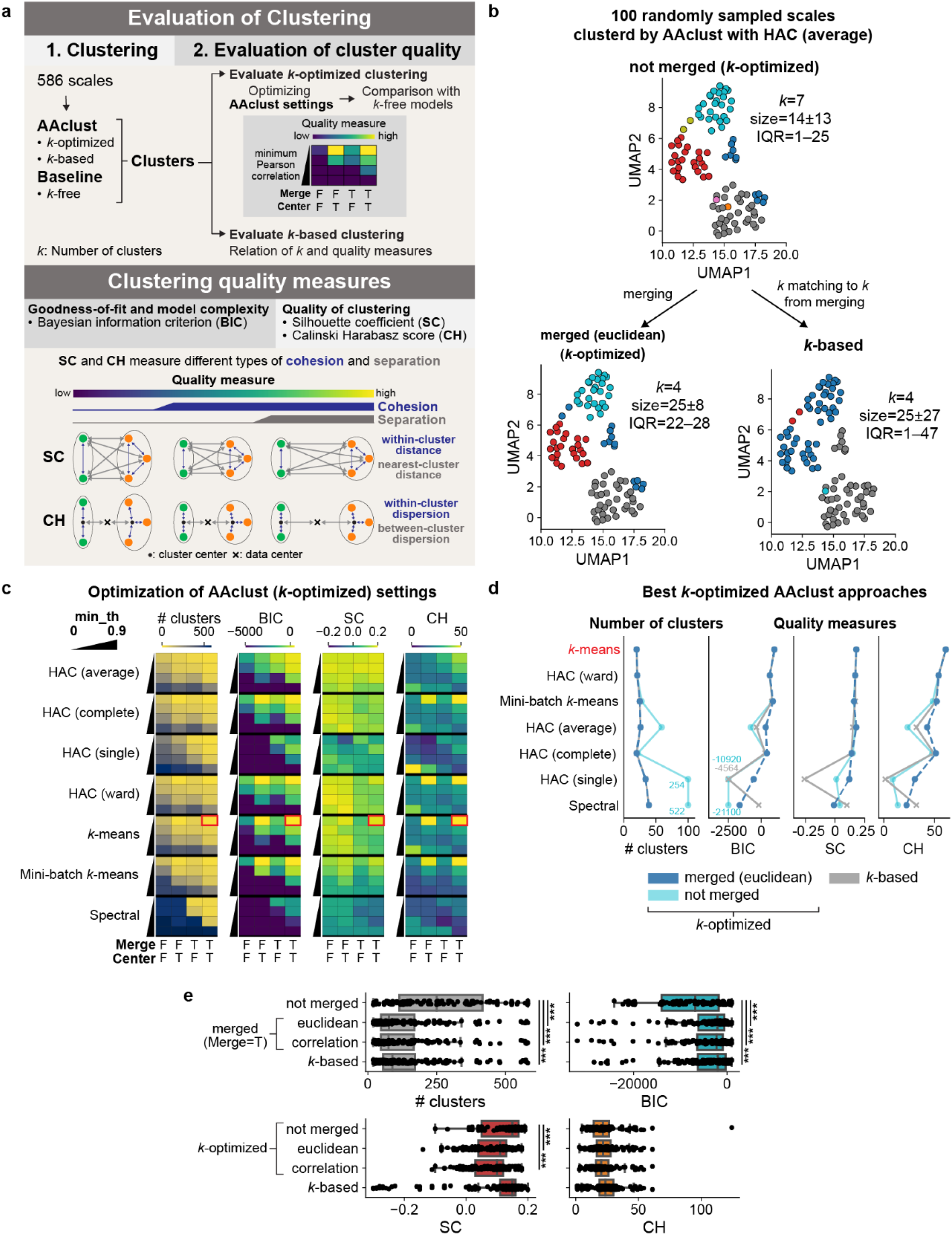
Evaluation of clustering by *k-*optimized AAclust approaches. **a**, AAclust clustering evaluation workflow and quality measures: Bayesian information criterion (BIC), silhouette coefficient (SC), and Calinski Harabasz score (CH), which gauge cluster cohesion and separation (see ‘3.1 Quality measures for clustering’). **b**, Effect of merging on clustering: AAclust (hierarchical agglomerative clustering (HAC) with average linkage) produces more uniform clusters in terms of size (noted by standard deviation and inter quantile range (IQR)) compared to *k*-based method. **c**, Optimization results for AAclust settings (cluster number and three quality measures) across seven clustering models. The minimum correlation threshold (*min_th*) results are shown for *min_th*=0, 0.3, 0.6, 0.9. Whether merging (Euclidean distance) was applied or not is indicated by T (True) and F (False), respectively; likewise, for minimum with-center (‘Center’=T) or within-cluster correlation (‘Center’=F). Red squares indicate best results achieved by *k-*means (*min_th*=0, merging with *min_cor*_*center*_). **d**, Clustering results for best-performing models from c (*k*-means in red), comparing merging, not merging, and *k-based* AAclust approaches. **e**, Overall model results, differentiating between merging methods, and cluster sizes matching to *k*-based approaches. AAclust setting differences were tested by a one-way ANOVA with Sidak post hoc test (\**P<*0.05,\*\**P<*0.01, \*\*\**P<*0.001).

**Fig. S2.**
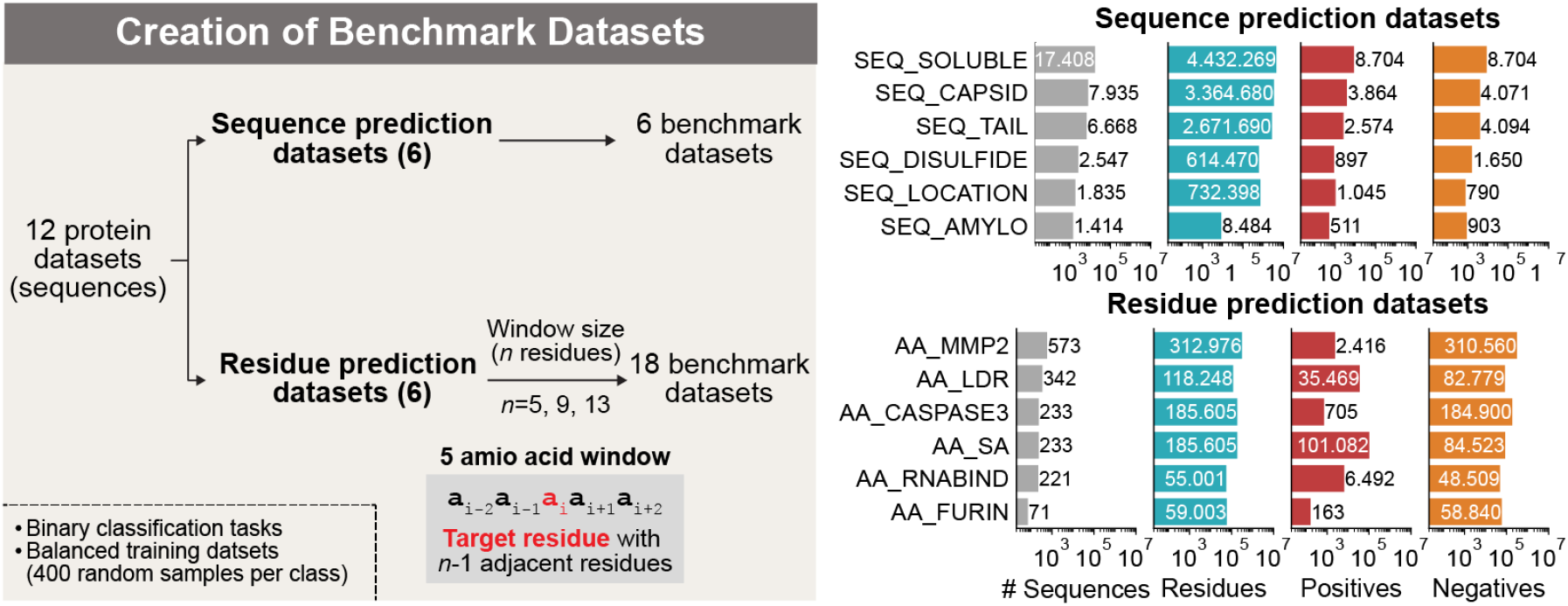
Creation of benchmark datasets. Workflow of benchmark dataset creation (left) and overview of the sequence and residue prediction datasets (right) used as machine learning benchmark datasets for scale set evaluation.

**Fig. S3.**
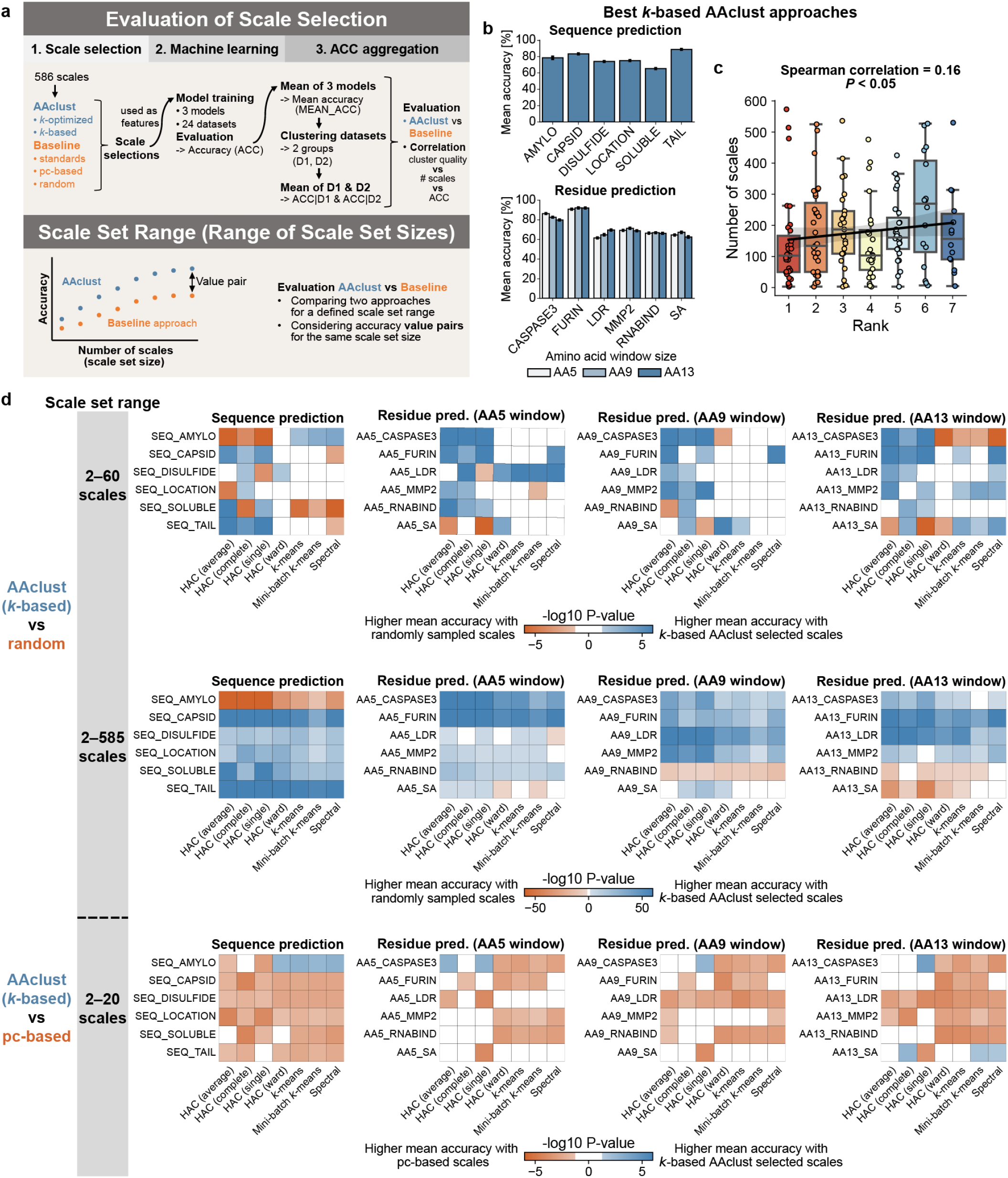
Evaluation of scale selection by *k*-based AAclust approaches. **a**, Workflow of AAclust scale selection evaluation and scheme of the scale set range used for the *k-*based AAclust analysis. **b**, Bar chart of mean accuracy (with standard deviation) per dataset for the best-performing models. Residue prediction datasets are grouped by amino acid window sizes (5, 9, 13). **c**, Number of scales per model rank, showing a positive Spearman’
ss correlation. **d**, AAclust (*k*-based) scale selection compared against baseline scale sets (random and pc-based) for seven clustering models. Ranges of 2–60 and 2–585 scales were used for comparison against random scales, and 2–20 for the pc-based scale set. AAclust results were tested against the corresponding baseline set using the paired Wilcoxon signed-rank test. Significantly better and worse AAclust performance is marked in blue and orange, respectively.

**Fig. S4.**
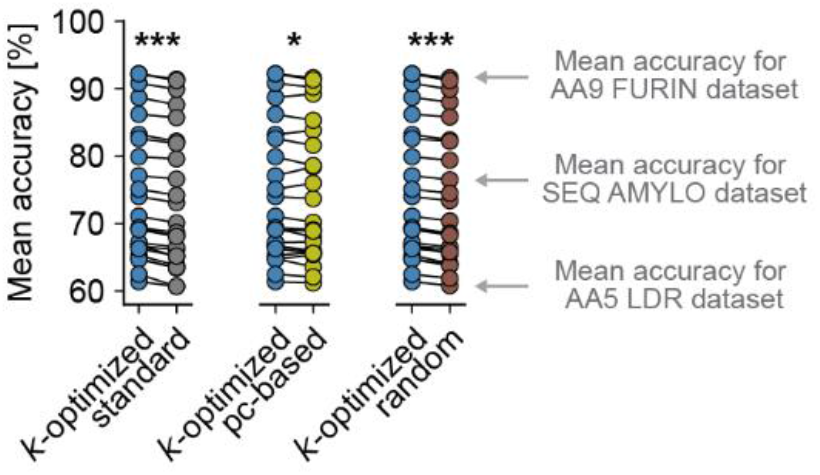
Comparison of *k*-optimized AAclust and baseline scale sets. Best-performing scale sets obtained *k*-optimized AAclust approaches are contrasted with best-performing baseline scale sets: ‘standard’ (gray), ‘pc-based’ (yellow), and ‘random’ (brown). For each of the 24 benchmark dataset, the scale sets achieving the highest prediction accuracy were considered. Differences were tested by paired Wilcoxon signed-rank test, Benjamin Hochberg correction (*P<0.05,**P<0.01, ***P<0.001). See **Fig. 1g** for comparison of *k*-based AAclust approaches against baseline scale sets.

**Fig. S5.**
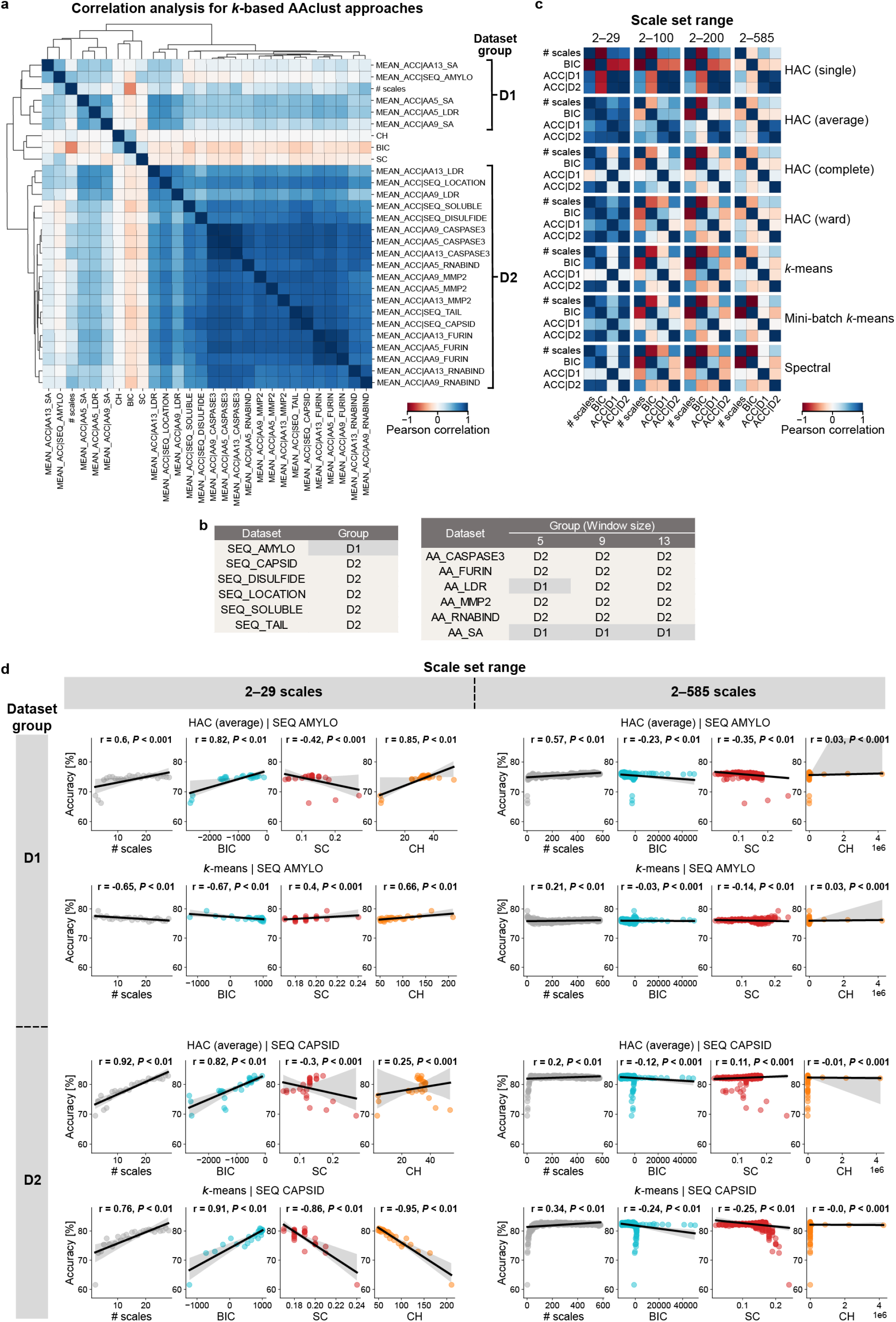
Dataset aggregation and correlation analysis. **a**, Hierarchical clustering of benchmark datasets utilizing Pearson correlation of average machine learning model performance (accuracy (ACC)) across seven AAclust approaches by varying the number of scales. **b**, Protein dataset grouping for sequence and residue prediction datasets. **c**, Nested heatmap showcasing Pearson correlation between average model performance for groups D1 and D2 (ACC|D1 and ACC|D2), the number of scales, and BIC, across seven clustering models and four scale set ranges. **d**, Examples of Pearson correlation for selected scale set ranges (2–29, 2–585), datasets (SEQ_AMYLO, SEQ CAPSID), and clustering models (HAC (average), *k-*means).

**Fig. S6.**
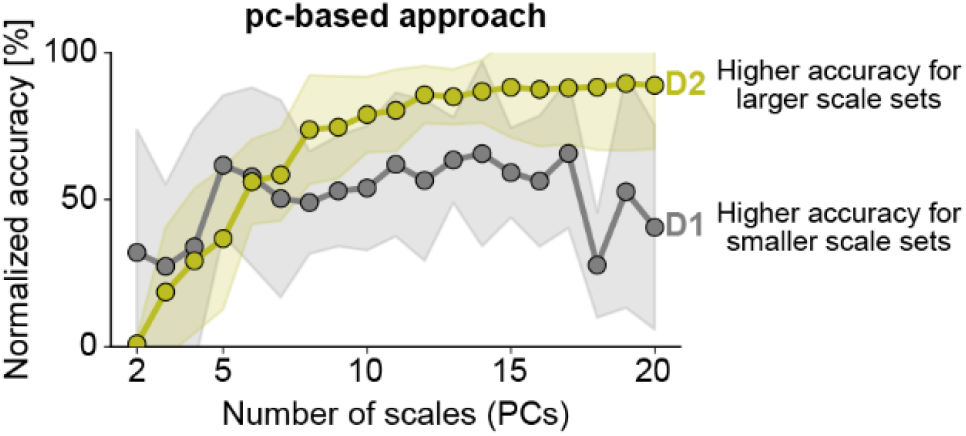
Evaluation of pc-based scale sets. Line plot showing min-max normalized accuracy for principal component (PC)-based scale sets averaged for dataset groups D1 (gray) and D2 (yellow).

**Fig. S7.**
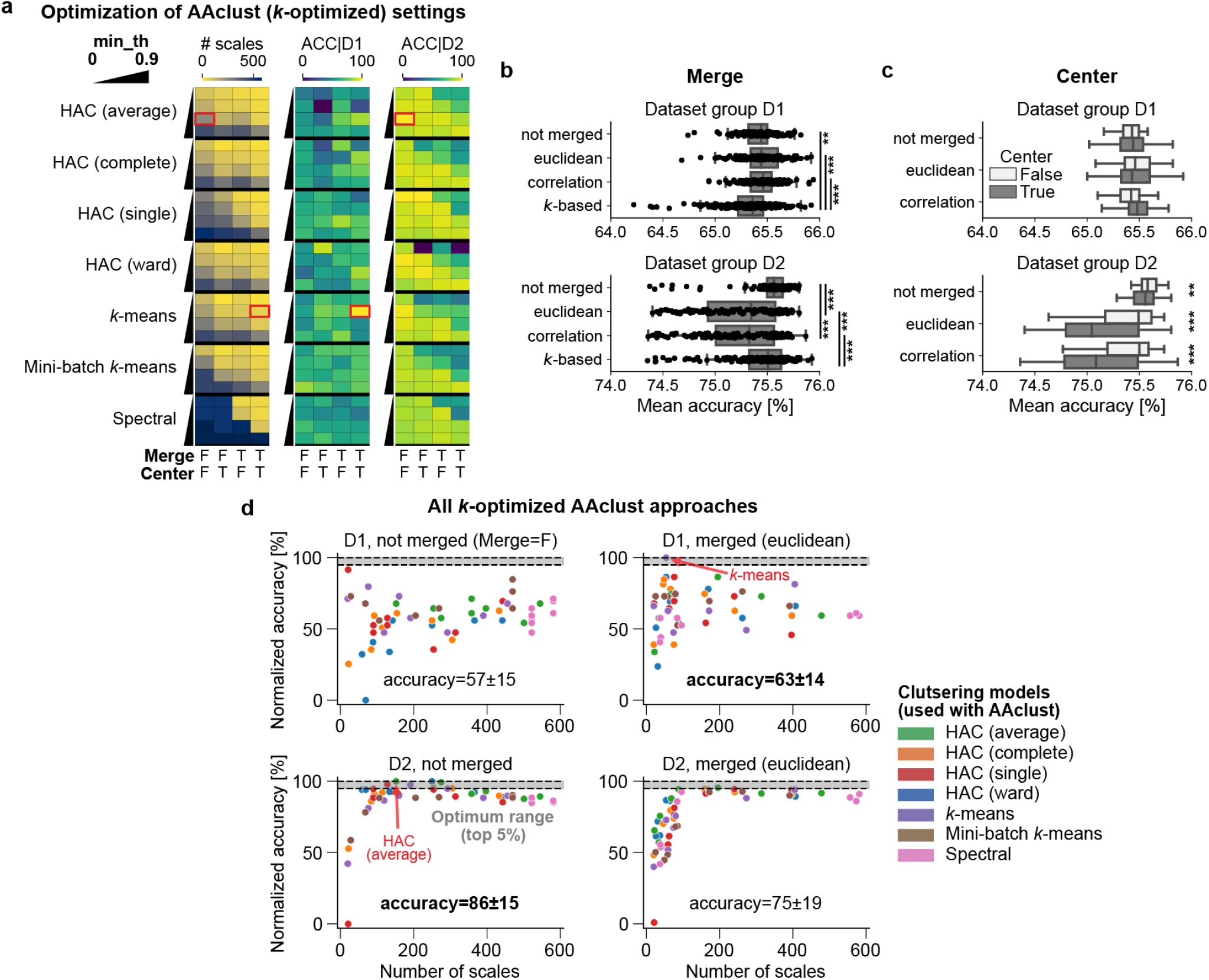
Evaluation of scale selection by *k-*optimized AAclust approaches. **a** Comparison of the number of scales and scale selection results, represented as aggregated accuracy for dataset groups D1 (ACC|D1) and D2 (ACC|D2), for AAclust settings across seven clustering models. Results are displayed for minimum correlation thresholds (*min_th*) of 0, 0.3, 0.6, 0.9. The use of merging (with Euclidean distance) is indicated as T (True) or F (False); similarly, the application of *min_th* is denoted for the cluster center (Center=T) or all cluster members (Center=F). **b-c**, Box plots showcasing mean accuracy across various AAclust settings—not merged, merged with Euclidean distance, merged with Pearson correlation, *k*-based clustering with matching number of clusters. Distinctions are made by D1 and D2 dataset groups in (b) and further by the ‘Center’ AAclust setting in (c). Statistical differences were tested by a one-way ANOVA with Sidak post hoc test in (b) and independent t-test with Bonferroni correction in (c) (\**P<*0.05,\*\**P<*0.01, \*\*\**P<*0.001). **d**, Min-max normalized accuracy plotted against the number of scales for seven clustering models used by AAclust. The optimum range, reflecting the top-5% performance (*i*.*e*., normalized accuracy >=95%), is highlighted in gray. The best-performing models are marked by red arrows (*k*-means for D1, HAC (average) for D2).

**Fig. S8.**
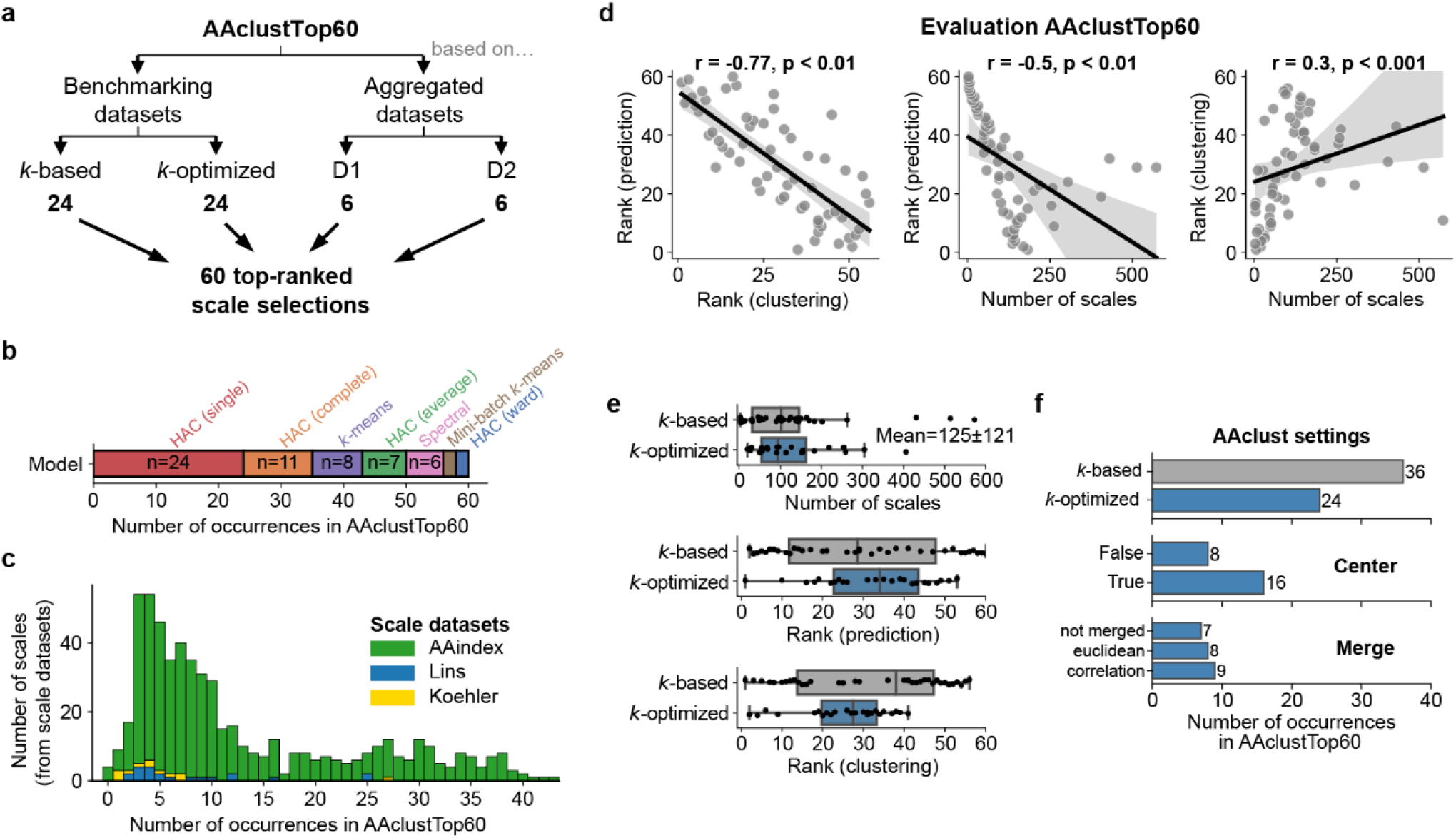
Evaluation of AAclustTop60. **a**, Collation of the best 60 scale sets, ‘AAclustTop60’, encompassing (a) 48 sets from the top-ranked *k*-based and *k*-optimized AAclust approaches for 24 benchmark datasets and (b) 12 sets derived from the top-ranked scale sets based on the aggregated prediction performance for dataset groups D1 and D2. **b**, Overview of clustering models used in AAclustTop60. **c**, Stacked histogram illustrating the frequency distribution of scales present in AAclustTop60, color-code indicating their origin. **d-f**, In-depth assessment of AAclustTop60. **d**, Scatterplots showing pairwise Pearson’s correlations of scale sets regarding their prediction performance-based ranking (average ranking over D1 and D2 accuracies), clustering-based ranking (average ranking for quality measures), and their number of scales. **e**, Boxplots comparing *k*-based and *k-*optimized AAclust approaches regarding parameters from (d). **f**, Barplots showing the number of occurrences in AAclustTop60 for *k*-based and *k*-optimized AAclust approaches and for AAclust optimization settings—namely the objective function (‘Center’) and the merging technique.

## Notes

### Competing Interest Statement

The authors have declared no competing interest.

